# Jaw size variation is associated with a novel craniofacial function for galanin receptor 2 in an adaptive radiation of pupfishes

**DOI:** 10.1101/2023.06.02.543513

**Authors:** M. Fernanda Palominos, Vanessa Muhl, Emilie J. Richards, Craig T. Miller, Christopher H. Martin

**Affiliations:** Department of Integrative Biology, University of California, Berkeley; Museum of Vertebrate Zoology, University of California, Berkeley; Department of Ecology, Evolution, and Behavior, University of Minnesota; Department of Molecular & Cell Biology, University of California, Berkeley

**Keywords:** gene expression, fluorescence in situ hybridization, galanin, trophic morphology, craniofacial divergence, evo devo, adaptive radiation

## Abstract

Understanding the genetic basis of novel adaptations in new species is a fundamental question in biology that also provides an opportunity to uncover new genes and regulatory networks with potential clinical relevance. Here we demonstrate a new role for *galr2* in vertebrate craniofacial development using an adaptive radiation of trophic specialist pupfishes endemic to San Salvador Island in the Bahamas. We confirmed the loss of a putative *Sry* transcription factor binding site in the upstream region of *galr2* in scale-eating pupfish and found significant spatial differences in *galr2* expression among pupfish species in Meckel’s cartilage and premaxilla using in situ hybridization chain reaction (HCR). We then experimentally demonstrated a novel function for Galr2 in craniofacial development and jaw elongation by exposing embryos to drugs that inhibit Galr2 activity. Galr2-inhibition reduced Meckel’s cartilage length and increased chondrocyte density in both trophic specialists but not in the generalist genetic background. We propose a mechanism for jaw elongation in scale-eaters based on the reduced expression of *galr2* due to the loss of a putative *Sry* binding site. Fewer Galr2 receptors in the scale-eater Meckel’s cartilage may result in their enlarged jaw lengths as adults by limiting opportunities for a postulated Galr2 agonist to bind to these receptors during development. Our findings illustrate the growing utility of linking candidate adaptive SNPs in non-model systems with highly divergent phenotypes to novel vertebrate gene functions.

## Introduction

Craniofacial developmental anomalies are the most common source of birth defects in humans, present in 1 out of 700 births (Hall et al., 2010; Meng et al., 2009; Palmer et al., 2016). While Mendelian craniofacial defects are well characterized (e.g. Treacher Collins Syndrome (Kadakia et al., 2014), Apert Syndrome (Wilkie et al., 1995), and Crouzon Syndrome (Helman et al., 2014; Reardon et al., 1994)), the developmental genetics of complex craniofacial defects, such as micrognathia, are poorly understood (Carlborg and Haley, 2004; Glazier, 2002; Hirschhorn and Daly, 2005; Hochheiser et al., 2011; Manolio et al., 2009; Monteiro and Podlaha, 2009; Weinberg et al., 2018). With the continued lowering costs of genomic sequencing and functional genetic tools, it is increasingly feasible to develop new genetic models for understanding human development and disease.

Current knowledge of craniofacial developmental genetics results mainly from mutagenic screens and forward genetic approaches in a few vertebrate model systems (primarily mice and zebrafish); however, these approaches are typically biased toward large Mendelian effects on easily detectable phenotypes, often missing the variation most relevant to common craniofacial birth defects in humans, such as small-effect variants, subtle differences in skeletal shape, and epistatic interactions among genes and the environment (Albertson et al., 2009; Powder and Albertson, 2016; Roberts et al., 2011). Conversely, complementary genome-wide association studies (GWAS) of craniofacial syndromes and normal-range facial variation in humans often reveal small-effect loci with no tractable possibilities for functional interrogation (Bonfante et al., 2021; Naqvi et al., 2022; Weinberg et al., 2018). Thus, understanding the genetic bases of naturally occurring, highly divergent adaptive phenotypes in novel systems that parallel human clinical variation, such as ‘evolutionary mutant’ models (Albertson et al., 2009; Concannon and Albertson, 2015; Powder and Albertson, 2016; Rohner, 2018), provides a powerful approach combining the tractable functional investigations possible in vertebrate model systems with genome-wide association scans of small-effect regulatory loci underlying natural craniofacial diversity. For example, novel insights into a broad diversity of human diseases have now arisen from studies of diverse species experiencing strong selective pressures in nature, including senescence (i.e., naked mole rat (Buffenstein, 2008); turquoise killifish (Platzer and Englert, 2016), and rockfish (Kolora et al., 2021)); eye degeneration, bone loss and enhanced lipogenesis (blind cavefishes (Gross et al., 2016; Hofmann et al., 2009; Protas et al., 2007; Lam et al. 2022)); body size (domesticated dogs (Sutter et al., 2007). In particular, the most remarkable diversity of vertebrate craniofacial morphology is represented in teleost fishes, often associated with their diverse and sometimes highly specialized modes of feeding (e.g., Liem, 1991; Otten, 1983; Albertson et al., 2003; Martin and Richards 2019; Konings et al., 2021; Evans et al. 2021).

Emerging fish model systems include the rapidly evolving East African and Cameroon cichlid radiations, in which a small number of genetic changes underlie immense morphological disparity in over 1,500 cichlid species (Brawand et al., 2014; Conith et al., 2019; Kocher et al., 2003; Martin, 2012; Martin, 2013; Martin and Genner, 2009; Navon et al., 2020) and the repeated parallel speciation of stickleback ecomorphs in glacial lakes (Chan et al., 2010; Erickson et al., 2016; Jones et al., 2012; Miller et al., 2007; Ishikawa et al. 2019). These systems provide excellent examples of leveraging naturally-occurring and highly divergent craniofacial phenotypes as ‘evolutionary mutants’ or ‘evolutionary forward genetics’ models to gain novel insights into the genetics of natural human craniofacial variation (Erickson et al., 2014; Glazer et al., 2015; J. Parsons et al., 2021; Kocher, 2004; Powder and Albertson, 2016; Roberts et al., 2009).

Here we demonstrate the utility of a an evolutionary radiation of *Cyprinodon* pupfishes for discovering and validating the craniofacial function of a new gene associated with jaw evolution and identified by QTL and GWAS analyses. Pupfishes offer some advantages over other evolutionary fish systems because they 1) rapidly evolved highly divergent and unique craniofacial phenotypes (Fig. 1) with minimal genetic differentiation among species (Hernandez et al., 2018; Lencer et al., 2016; Lencer et al., 2017; Martin and Wainwright, 2011; Martin and Wainwright, 2013a; Martin et al., 2019; St John et al., 2020a; St John et al., 2020b), 2) speciated in the face of ongoing gene flow resulting in very few highly differentiated genomic regions associated with species-specific craniofacial traits (Martin and Feinstein 2014; Martin et al., 2017; McGirr and Martin, 2016; McGirr and Martin, 2021; Richards and Martin, 2017; Richards and Martin, 2022; Richards et al., 2021), and 3) are highly amenable to laboratory rearing and imaging due to their high fecundity, daily egg production, and egg transparency comparable to zebrafish (Martin and Gould, 2020; Martin and Wainwright, 2013b). This radiation contains the widespread algae-eating generalist pupfish, *Cyprinodon variegatus* (Fig. 1a), which is broadly distributed across the Caribbean and North American Atlantic coast and occurs in sympatry with two microendemic trophic specialist species found only in the hypersaline lakes of San Salvador Island (SSI), Bahamas. Each trophic specialist displays highly divergent behavior, pigmentation, and craniofacial morphology: the molluscivore *C. brontotheroides*, has a novel nasal protrusion that is a skeletal extension of the maxilla and foreshortened robust oral jaw (Fig. 1b); and the scale-eater, *C. desquamator*, exhibits two-fold larger oral jaws and overall brachycephalic features (Fig. 1c) (Martin, 2016; Martin and Wainwright, 2011; Martin and Wainwright, 2013a). There is also a fourth intermediate scale-eating ecotype in some lakes (Richards et al. 2022).

**Figure 1.**
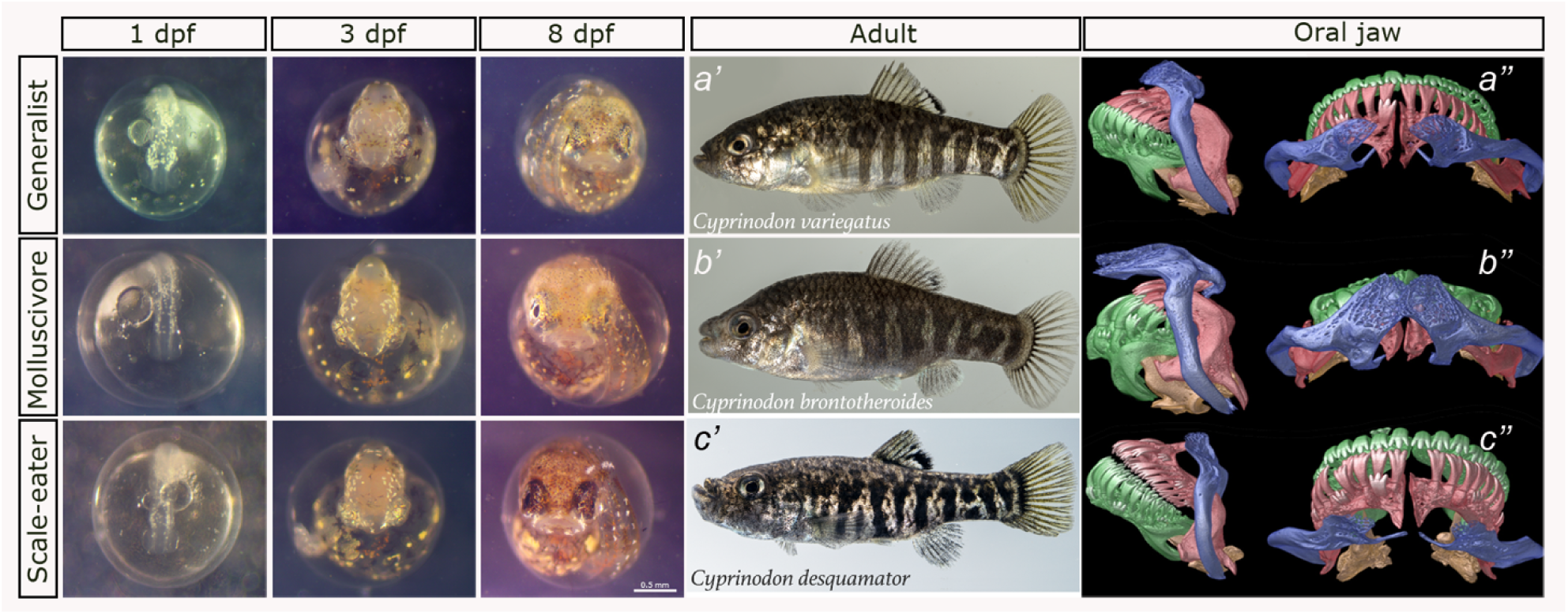
Divergent craniofacial morphology and development of the San Salvador Island *Cyprinodon* pupfish radiation. *C. variegatus* (first row) is a trophic generalist distributed across the western Atlantic and Caribbean; *C. brontotheroides* (second row) is a molluscivore, and *C. desquamator* (third row) is a scale-eater, both endemic to the hypersaline lakes of San Salvador Island (SSI), Bahamas. Left panel: development at 1-, 3-, and 8-days post-fertilization (dpf). Middle panel: lab-reared adult female pupfishes of each species. Right panel: Lateral and dorsal views of µCT scans of craniofacial morphology (modified from Hernández et al., 2017). The maxilla is colored in blue, premaxilla in red, dentary in green, articular in orange.

Previous genomic and transcriptomic work on the SSI pupfishes identified dozens of new candidate craniofacial genes never previously characterized as craniofacial or directly investigated in other systems (McGirr and Martin 2016, 2019, 2020; Richards et al. 2021, 2022; Richards and Martin 2017; Patton et al. 2021; St. John et al. 2021). One of the most promising candidates was galanin receptor 2a, the second receptor type for galanin. Using a genome-wide association (GWA) test across 202 individuals from the SSI radiation and outgroup populations, we previously found an association of the regulatory region of *galr2a* with lower jaw length, confirming an earlier pilot study that found the *galr2a* region to be among the top five strongest associations with lower jaw length, containing highly differentiated SNPs between trophic specialist species (Richards et al. 2021; McGirr and Martin 2017). An analysis of hard selective sweeps using both site frequency spectrum (SweeD) and linkage disequilibrium (Omegle) based summary statistics additionally found evidence of a putative adaptive allele in the 20 kb regulatory region upstream of *galr2* that swept to fixation in the scale-eater *C. desquamator* population on SSI 696-1,008 (95% credible interval) years ago, potentially providing a pivotal stage in adaptation to scale-eating (Richards et al. 2021). Furthermore, an independent quantitative trait loci (QTL) mapping study in F2 intercross hybrids between scale-eater and molluscivore parents found a significant QTL on linkage group 15 containing *galr2a* that accounted for 15.3% of the phenotypic variance in premaxilla length (*n* = 178; Martin et al. 2016), with a positive effect on jaw length in the scale-eater genotype. Similarly, a second independent study of F2 hybrid intercross from a second lake found evidence of a QTL in this region explaining 8% of the phenotypic variance in the length of the coronoid process on the articular bone of the lower jaw (jaw closing in-lever; *n* = 227; St. John et al. 2021). The combined strength of evidence for a role of *galr2a* in craniofacial development across independent analyses of GWA, QTL, selective sweeps, and genetic differentiation between species indicated that this was one of our highest priority candidates for functional studies.

In humans, *galr2* is abundantly expressed within the hypothalamus and hippocampus of the central nervous system and in the heart, kidney, liver, colon, and small intestine, and has genetic associations with epilepsy and Alzheimer’s (Li et al., 2013; Ma et al., 2018). Classified as an orexigenic (appetite stimulant) gene, *galr2* is expressed in the human hypothalamus but also in the ventral telencephalon of larval and adult zebrafish (Ahi et al., 2019; Li et al., 2013; Kim et al., 2016). Although a role of *galr2* in craniofacial development has not previously been reported in the literature, its ligand, the neuropeptide galanin (GAL), is highly expressed in bones from early to post-embryonic development (Xu et al., 1996, Jones et al., 2009), with demonstrated effects on bone mass (Idelevich et al., 2018; Idelevich et al., 2019), muscle contraction (Kakuyama et al., 1997. EJP), and periodontal regeneration (Ma et al. 2021).

Here we used Sanger sequencing to confirm two highly differentiated SNPs between SSI specialist species detected in our previous genomic studies affecting two predicted transcription factor binding sites in the *galr2* regulatory region, characterized the divergent craniofacial expression of *galr2* across all three SSI pupfish species using *in situ* hybridization chain reaction (HCR) (Choi et al., 2018; Ibarra-Padilla-García et al., 2020) at two key developmental timepoints, and demonstrated that treatment with two different Galr2 receptor inhibitors reduced Meckel’s cartilage length and increased chondrocyte density dependent on the species’ genetic backgrounds.

## Results

### Two highly differentiated SNPs are associated with *galr2a* transcription factor binding sites

Previous whole genome resequencing of over one hundred San Salvador Island (SSI) pupfishes identified only two highly differentiated single nucleotide polymorphisms (SNP) within the 20 kb regulatory region of *galr2a* between trophic specialists across different lake populations on SSI (Richards et al. 2021). We designed primers (Table S5) and used Sanger sequencing to further genotype these SNPs in a large panel of wild-caught specialists from six lake populations. We confirmed the presence of a transversion from G to A approximately 11 kb upstream of the *galr2a* transcription starting site (TSS; Fig. 2a., S1; Table S1), which changes a predicted CA**G**CAA *Elf1/Erg* transcription factor binding site (TFBS) to a predicted **A**GGGASW *Elf5* TFBS at this locus (using the Multiple Expectation maximizations for Motif Elicitation (MEME) server and the motif database scanning algorithm TOMTOM (Bailey et al., 2009)). This transversion was observed in 86.3% of scale-eaters across four lake populations (*n* = 51) versus 20% of molluscivores across six lake populations on SSI (*n* = 30) (Fig. 2a., S1).

**Figure 2.**
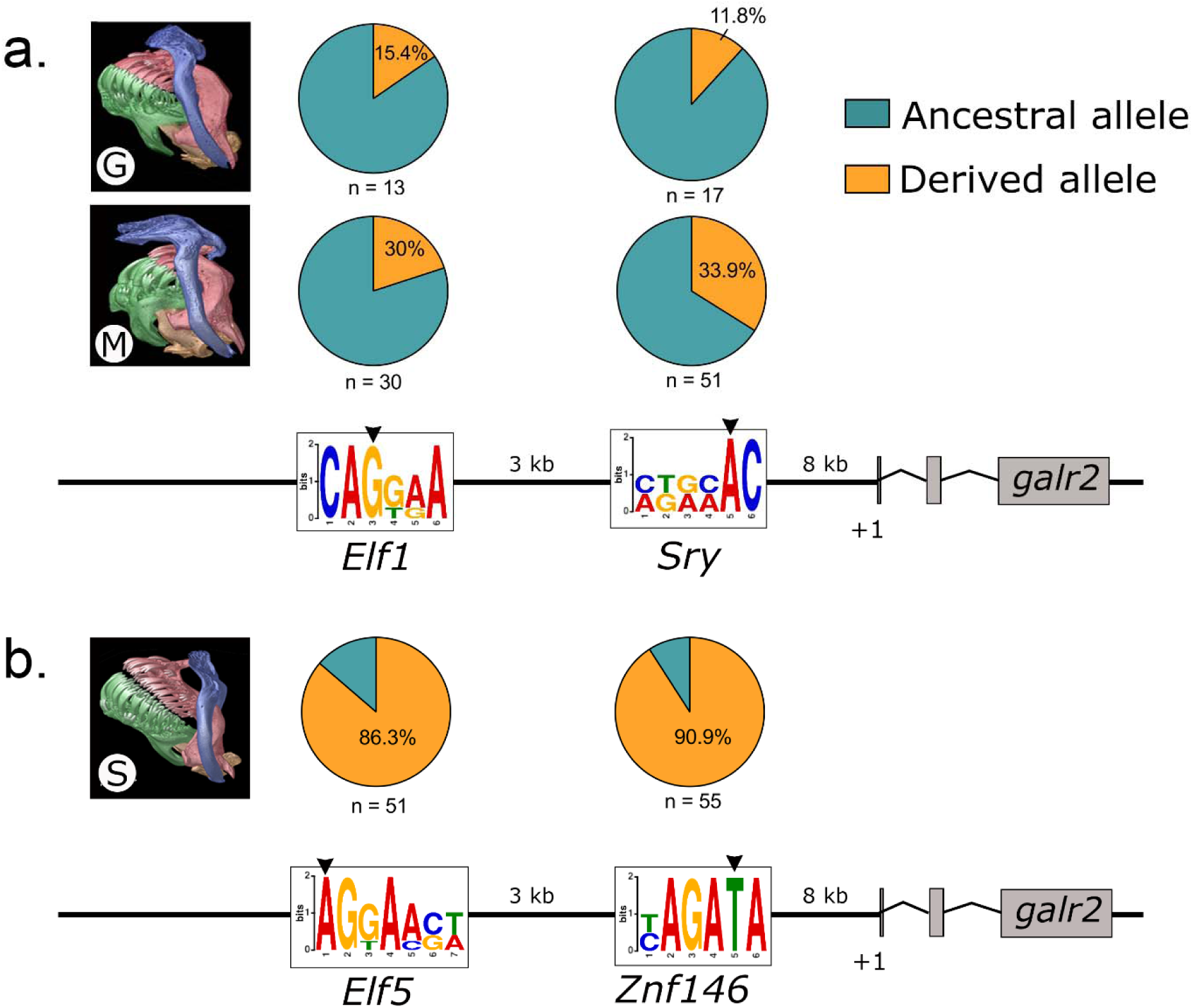
The two most differentiated single-nucleotide polymorphisms (SNP) between trophic specialists lie within the 20kb regulatory region of *galr2a*. **a.** Putative ancestral TFBS for *Elf1/Erg* and *Sry*, respectively, in the generalist (G), *Cyprinodon variegatus* and the molluscivore (M) *Cyprinodon brontotheroides*. **B.** Derived changes in predicted TFBS to *Elf5* and *Znf146* in the scale-eater (S) *Cyprinodon desquamator*. Pie charts indicate frequencies of ancestral (teal) and derived (orange) alleles at each locus. +1 represents the transcription starting site of *galr2a* (Galanin receptor 2a). µCT scans of craniofacial morphology are modified from Hernández et al., 2017. Sequence logos of the Position Weight Matrixes (PWM) identified using MEME and TOMTOM.

Using Sanger sequencing, we genotyped a second transversion from C to T approximately 8 kb upstream of *galr2a* TSS, which changes the predicted AGA**C**AA *Sry* TFBS to a predicted YAGA**T**A *Znf146* TFBS (Fig. 2b., S2; Table S1). This transversion was observed in 90.9% of scale-eaters across six lake populations (*n* = 55) versus 33.9% of molluscivores across six lake populations on SSI (*n* = 53) (Figure S2). Notably, all scale-eaters sampled from Crescent Pond (CRP) contained the T transversion (*n* = 30) (Figure S2). The predicted TFBS changes in the *galr2a* cis-regulatory region across pupfishes, combined with a previous genetic mapping study that found a significant QTL in this region explaining 15% of phenotypic variance in oral jaw size (Martin et al. 2017), suggests that different spatial or temporal *galr2a* expression may underlie some of the craniofacial divergence in SSI pupfishes. Previous studies of allele-specific expression in SSI pupfishes were inconclusive due to lack of heterozygous sites in the *galr2a* transcripts (McGirr and Martin 2021).

### Different spatiotemporal patterns of *galr2a* expression in craniofacial tissues

To determine if spatial or temporal changes in *galr2a* expression underlie SSI craniofacial divergence, we assayed *galr2a* expression in 2 dpf embryos and 8 dpf larvae from two independent lake populations for each of the three SSI species using hybridization chain reaction (HCR) in-situ hybridization for *galr2a* and *tropomyosin 3b* (*tpm3b*), a component of thin filaments of myofibrils expressed in fish skeletal muscles (Dube et al., 2016), to visualize jaw and other cranial muscles. We also tested *galr2a* expression using orthogonal amplifiers labeled with three distinct fluorophores (see Methods for details) to ensure reliable detection of *galr2a* transcripts *in situ* across species and developmental stages (Supp. Video S1).

At 2 dpf, *galr2a* expression was detected in broad regions of the central nervous system (e.g. the posterior tectum and medial longitudinal fasciculus) of specialists (Fig. 3a,c) and generalists, with apparent higher expression in both specialist species than in generalists. We also observed an apparent increase of *galr2a* expression anterior to the first pharyngeal arches in the molluscivore and generalist pupfishes relative to the scale-eaters.

**Figure 3.**
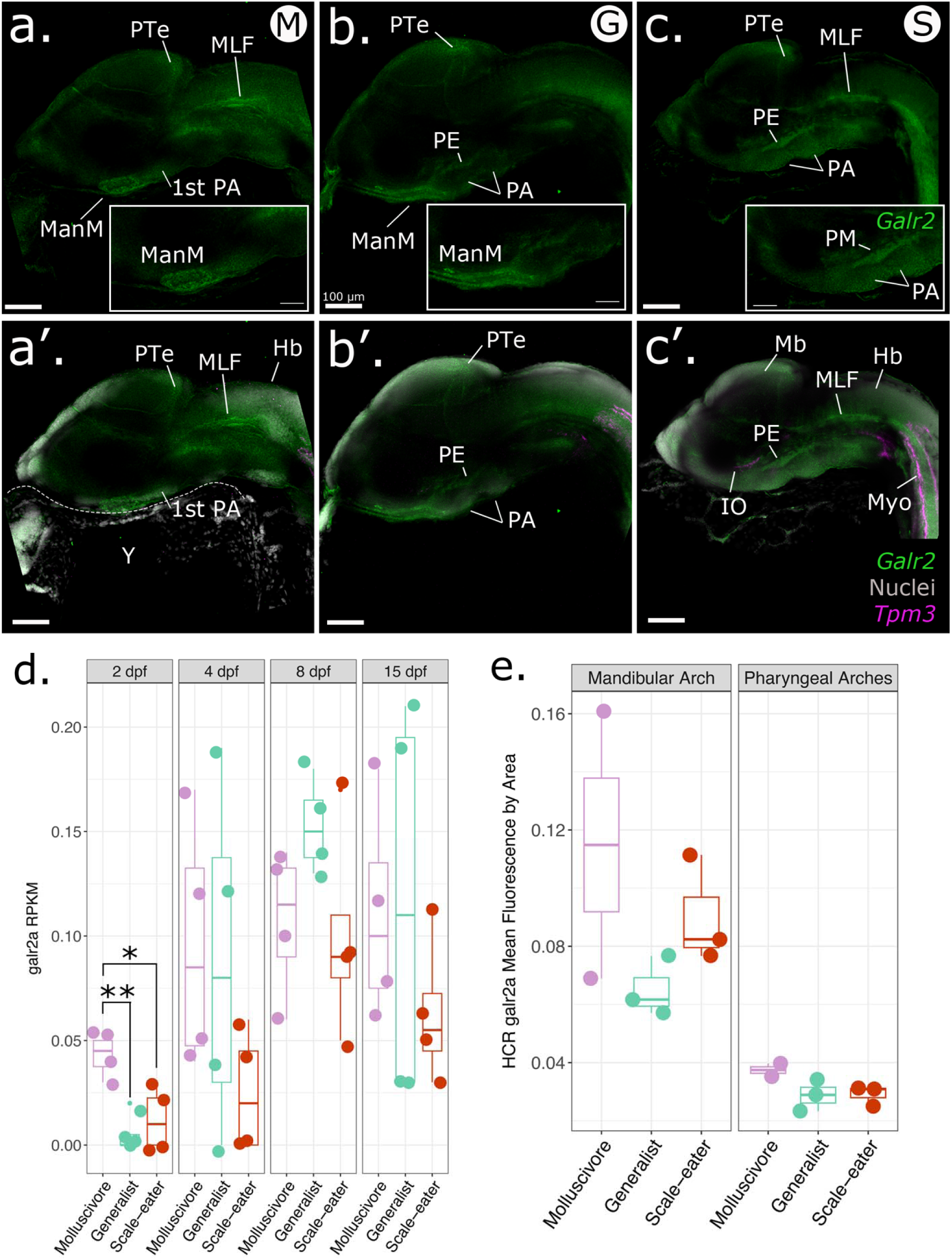
Spatiotemporal patterns of *galr2a* expression differ among SSI species at 2 dpf. Representative images of *Galr2a* (green) and *Tpm3b* (magenta) expression revealed by hybridization chain reaction (HCR) and DAPI nuclear staining (gray) in the: **a.** molluscivore, *C. brontotheroides*; **b.** generalist, *C. variegatus*; and **c.** scale-eater, *C. desquamator*. **d.** *Galr2a* reads per kilobase of transcript per million reads (RPKM) from an existing mRNAseq study sampling the entire head of each species at four developmental stages (Lencer and McCune, 2020). Galr2a showed significantly higher expression in the molluscivore relative to generalists (P = 0.003, Tukey’s HSD test) and scale-eaters (P = 0.014, Tukey’s HSD test) at 2 post fertilization (dpf). **e.** Mean fluorescence per area for *galr2a* quantified at 2 dpf in all three species shows similarly elevated levels of expression in the molluscivore for the mandibular and pharyngeal arches. Images shown are representative Z-stack maximum projection images of 30 optical sections taken every 3 µm from whole-mounted embryos imaged using a Zeiss LSM880 laser confocal microscope. PA: pharyngeal arches, PE: pharyngeal endoderm, ManM: mandibular mesenchyme, MLF: medial longitudinal fasciculus, PTe: posterior tectum, Di: diencephalon; Mb: midbrain, Hb: hindbrain, Y: yolk, Myo: somitic myofibers, IO: inferior oblique muscle.

Using 3D reconstruction and volume rendering analysis of HCR data for whole-mounted pupfishes at hatching time (8 dpf), we found that *galr2a* expression domain was expanded in the jaws of the molluscivores than in the generalists (Fig. 4; *P* = 0.03, Tukey’s HSD), consistent with either greater tissue volume or an increased gene expression domain. In contrast, *galr2a* showed no differences in expression volume among species in the brain and head (Fig. 4, Table S3). G*alr2a* expression was detected in the Meckel’s and palatoquadrate cartilages in all SSI pupfishes, in the premaxilla of the generalist and the scale-eater SSI specialist (Fig. 4 b-b’, c-c’), the maxilla of the molluscivore SSI specialist (Fig. 4 a-a’), and in the intermandibular muscles of the scale-eaters (Fig. 4 c’).

**Figure 4.**
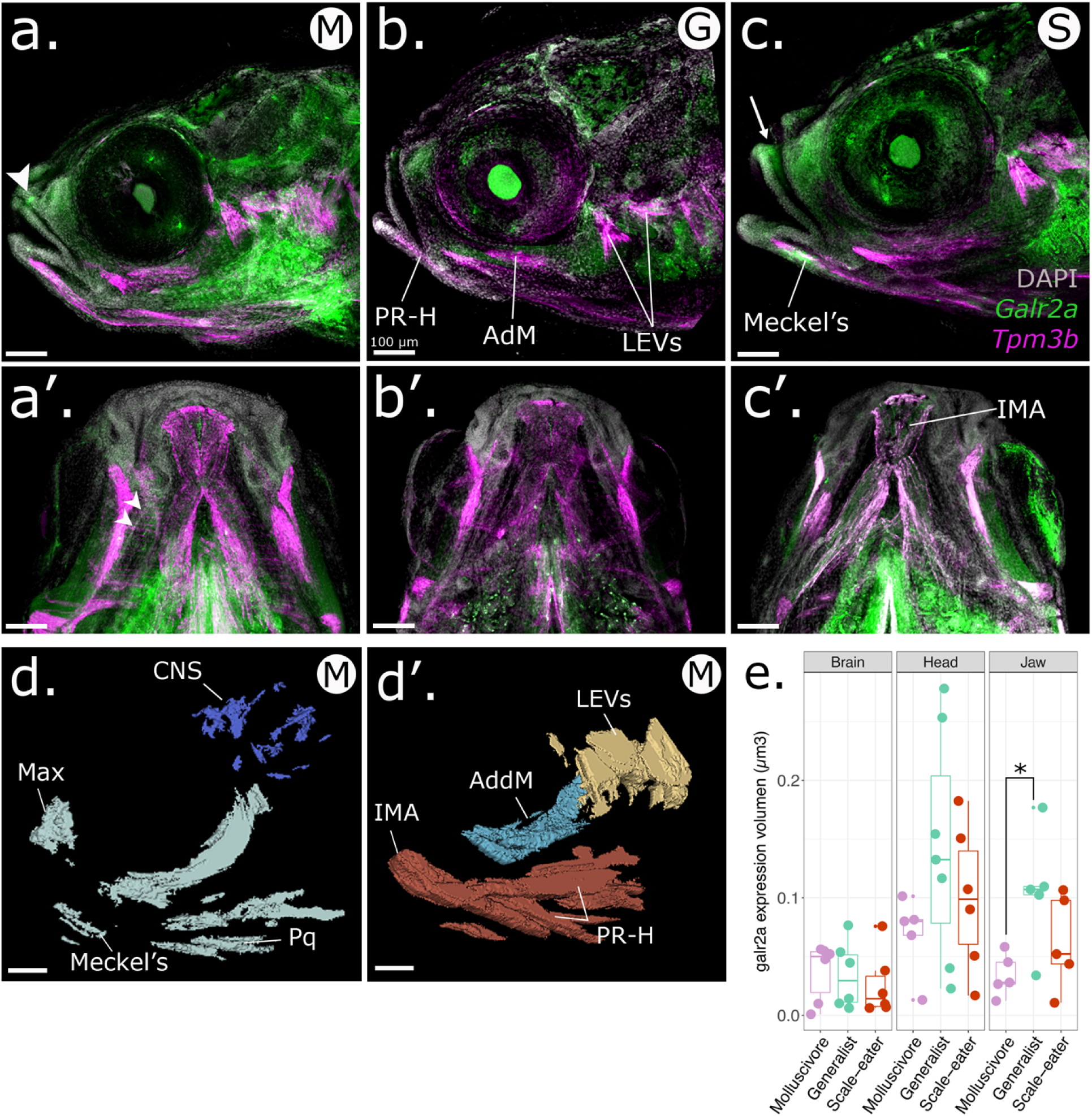
Spatiotemporal patterns of *galr2a* expression differ in molluscivores at 8 dpf. Representative images of *Galr2a* (green) and *Tpm3b* (magenta) expression revealed by hybridization chain reaction (HCR) and DAPI nuclear staining (gray). **a.** Lateral view of an 8 dpf molluscivore (M in white circle), **a’.** ventral view. **b.** Lateral view of an 8 dpf generalist (G in white circle), **b’.** ventral view. **c.** Lateral view of an 8 dpf scale-eater (S in white circle), **c’.** ventral view. The arrowhead in **b.** and the arrow in **c.** point to high expression of *galr2a* in the maxilla and premaxilla of the molluscivore and scale-eater specialists, respectively. White arrowheads in **a’.** point to high expression of *galr2a* in the palatoquadrate of the molluscivore. *Galr2a* expression in the IMA was found only in the scale-eaters (**c’**). Images shown here are representative Z-stack maximum projection images of 30 optical sections taken every 15 µm from whole-mounted larvae imaged using a Zeiss LSM880 laser confocal microscope. **d-d’.** 3D-Slicer view of the expression volume of *galr2a* (**d.**) and *tpm3b* (**d’.**) in an 8 dpf molluscivore pupfish larvae (*C. brontotheroides*). **e.** Volume of *galr2a* expression in the brain (left panel), whole head (middle panel), and jaw (right panel) across SSI pupfishes. CNS: central nervous system, Max: maxilla. LEVs: levator arcus palatini and operculi, AdM: adductor mandibular. IMA: intermandibular anterior. Total volume rendered: 450 µm. All scale bars: 100 µm.

At the subcellular level, *galr2a* mRNA was detected in the cytoplasm of chondrocytes at the Meckel’s symphysis (Fig. 5a). In contrast, on the distal edge of the Meckel’s cartilage expression was detected only in the cytoplasm of the elongated chondrocytes (Fig. 5a’). Towards the most posterior region of the Meckel’s cartilage closest to the palatoquadrate cartilage, we observed *galr2a* expression in the cells surrounding the jaw joint (Fig. 5 b, c, d; Supplementary Video S1). We conclude that g*alr2a* was significantly differentially expressed in specific and distinct craniofacial tissues in the specialists at hatching time, suggesting an important role for craniofacial divergence in the SSI radiation.

**Figure 5.**
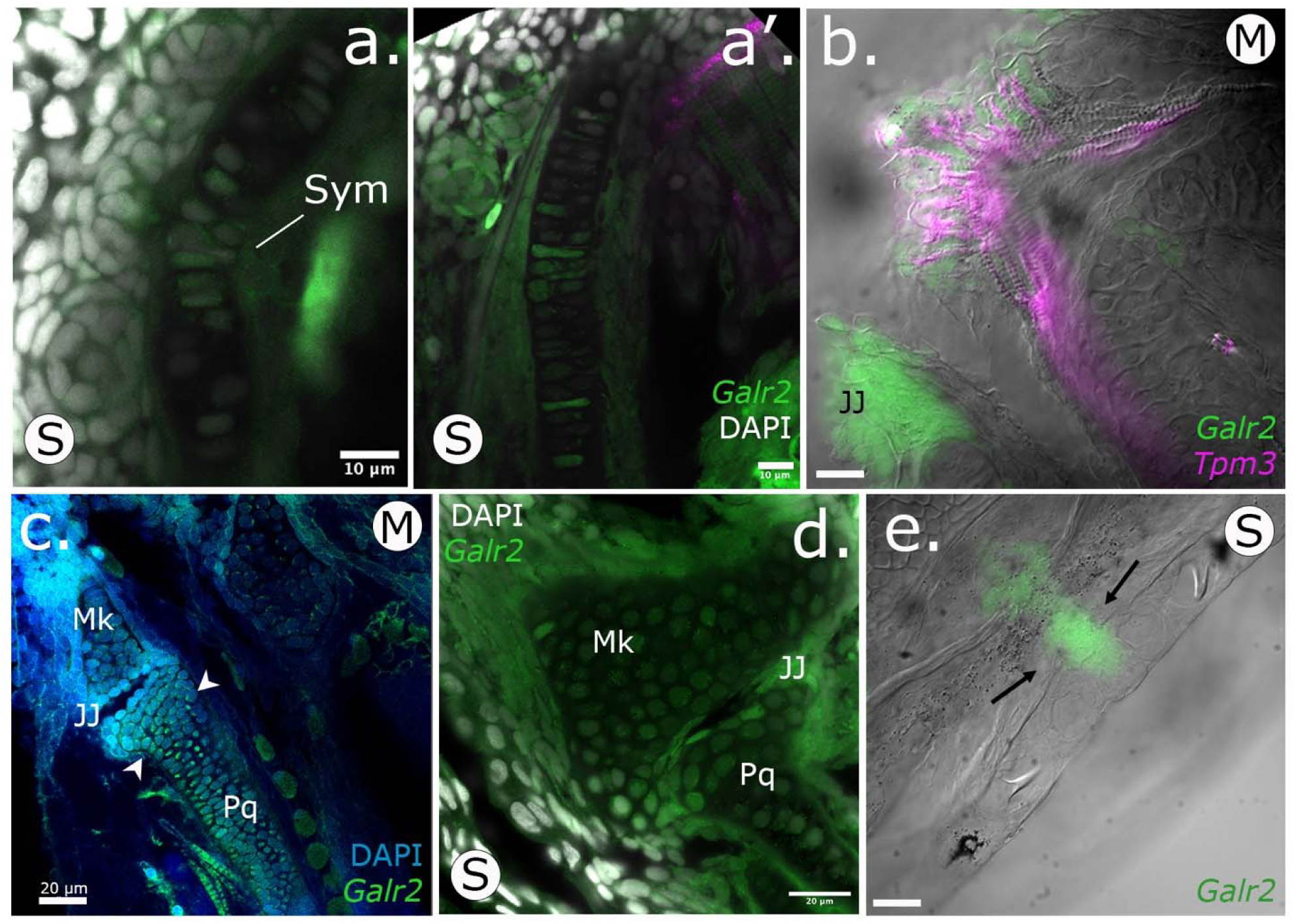
*Galr2a* is expressed in the chondrocytes of the Meckel’s and palatoquadrate cartilages at hatching (8 dpf) in SSI pupfishes. HCRs showing tissue expression of *Galr2a* in green, and the muscle marker Tropomyosin 3b, (*Tpm3b*), in magenta. Nuclei are stained with DAPI (gray). **a-a’.** Scale-eater *galr2a* expression in the chondrocytes of the Meckel’s symphysis (Sym) and lateral side. **b.** Ventral view of a molluscivore jaw at 8 dpf showing the expression of *Tpm3b* in the intermandibular muscles of the lower jaw. *Galr2a* is expressed in the jaw joint and the most anterior region of the lower jaw. **c-d.** *Galr2a* is expressed in the chondrocytes of the Meckel’s and palatoquadrate cartilages and around the jaw joint. **e.** *Galr2a* expression in the premaxilla of a scale-eater. Unlabeled scale-bars: 10 µm.

We further tested for quantitative differences *in galr2a* expression at 2 dpf and 8 dpf by quantifying transcript counts from the only existing RNAseq data set on SSI pupfishes heads during development (Lencer and McCune, 2020). Only *galr2a*, but not *galr1a, galr1b, galr2a, galr2b,* or *galanin*, showed significantly higher mRNA expression in the molluscivores relative to generalists (P = 0.003, Tukey’s HSD test) and scale-eaters (P = 0.014, Tukey’s HSD test) at 2 dpf (Table S2), whereas *galr2a* showed overall similar levels of expression in the head from 4 to 15 dpf in all three species (Table S2).

By using *Tpm3b* expression to label muscle cells, we observed at 2 dpf that *Tpm3b* expression was detected in somitic myofibers in all SSI pupfishes (Fig. 3a’’-c’’). However, only in the scale-eaters, *Tpm3b* expression was detected in the eye’s inferior oblique muscle primordial cells (Fig. 3c’’). At hatching time, *Tpm3b* expression was detected in all larval head muscles (Fig. 4a-a’, b-b’, c-c’; 5b). The scale-eater and molluscivore *Tpm3b* expression volume was significantly larger than in the generalists (Table S3; P < 0.05; 1-way ANOVA, Tukey’s HSD test).

### Chemical inhibition of Galr2 receptors affects Meckel’s cartilage length and chondrocyte density

To study the effect of Galr2 in craniofacial development, we inhibited the endogenous activation of all four known galanin receptors in teleost fishes (*Galr1a* and b, *Galr2a* and b; Butler et al. 2020, Li et al. 2013) using M35 (Innopep, Inc.), a synthetic peptide antagonist of Galr1+2 galanin receptors (Wiesenfeld-Hallin et al., 1993), and M871 (Abcam), a Galr2-specific synthetic peptide antagonist (Sollenberg et al., 2006). Embryos of all three species from two different lake populations were exposed from stages 24-25 (2 dpf, with the appearance of the first pharyngeal arches) until hatching at stages 32-33 (8 dpf, Fig. 6a).

**Figure 6.**
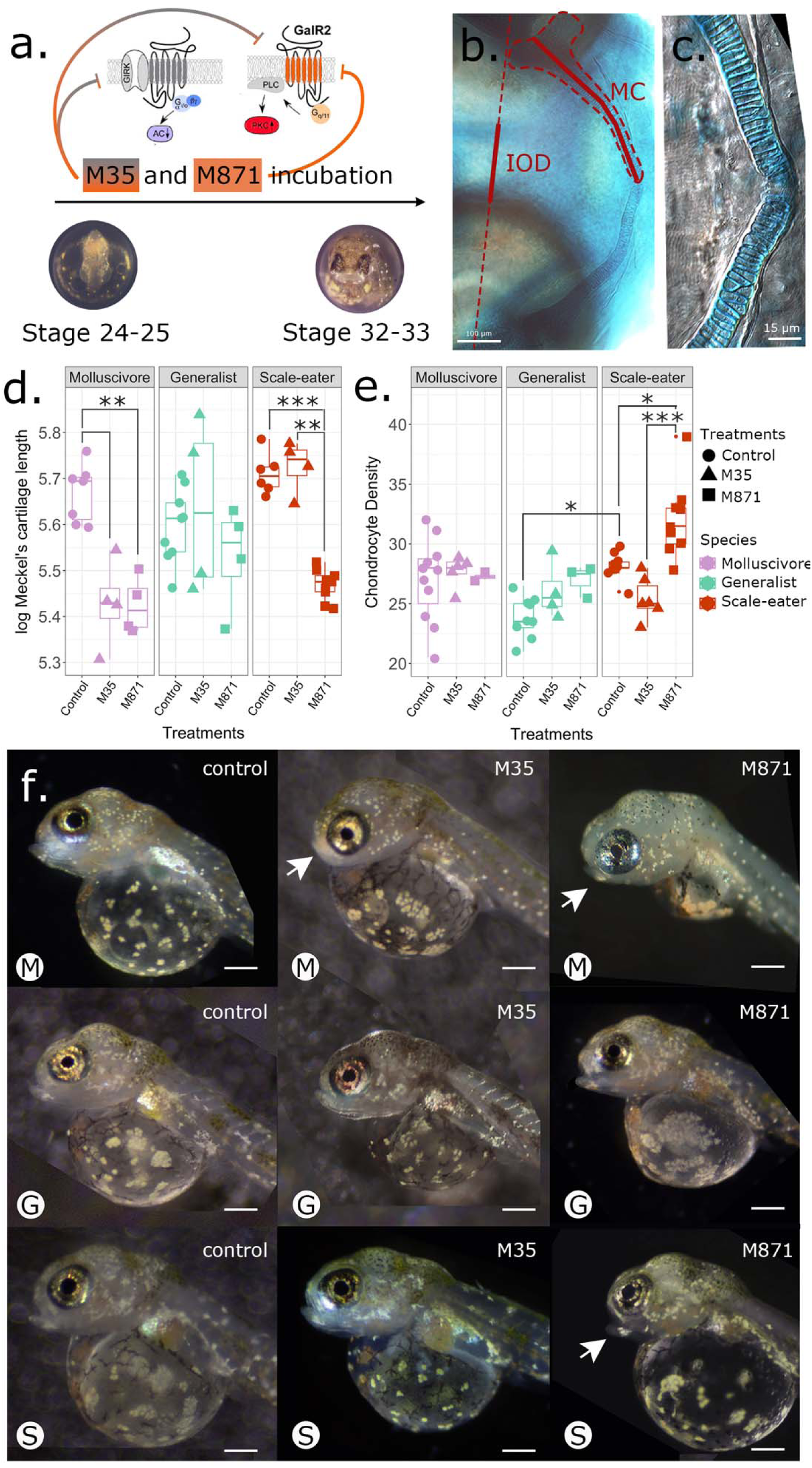
Galr2 receptor inhibition reduced Meckel’s cartilage length in both specialists and increased chondrocyte density in scale-eaters. **a.** Galr2 and Galr1+2 inhibition protocol during pupfish development. *C. desquamator* embryos are depicted. **b.** Alcian blue cartilage staining at Stage 32-33. IOD: interocular distance, MC: Meckel’s cartilage. The MC length and IOD are shown in red. **c.** Chondrocytes at the symphysis. **d.** Changes in the log-transformed Meckel’s length and the log-transformed interocular distance at stage 32-33 across species and treatments. **e.** Chondrocyte numbers at the symphysis across species and treatments. \**P* < 0.05, \*\**P* < 0.01, \*\*\**P* < 0.001 Tukey’s HSD test. **f.** Representative control and treated larvae of each species at stage 32-33. Arrows point to abnormal lower jaws. M: molluscivores, G: generalist, S: scale-eaters. Scale-bars = 0.4 mm. Galr1 and Galr2 cartoons were modified from Lang et al., 2007.

At approximate hatching time (8 dpf), both trophic specialist species raised under laboratory common garden conditions exhibited increased Meckel’s cartilage length (Fig. 6d), consistent with the longer jaws of adult scale-eaters and more robust jaws of the molluscivore relative to more gracile jaws of the generalist (Martin and Wainwright 2013). We found that exposure to M871, the Galr2-specific antagonist, significantly reduced the length of the Meckel’s cartilage in both specialists relative to the generalists (Fig. 6d.; P = 7.89 x 10^-5^ for scale-eaters; P = 0.001 for molluscivores; 2-way ANOVA, Tukey’s HSD test) while the interocular distance remained unchanged between control and treated larvae (Fig. S4). In contrast, M35 (Galr1 and Galr2 antagonist) only significantly reduced Meckel’s length in the molluscivore (Fig. 6c). Meckel’s length of generalists was unaffected by exposure to M35 or M871 (Fig. 6d).

To understand the cellular effect of Galr1 and 2 antagonists on Meckel’s length across species, we quantified the number of chondrocytes 100 µm from the symphysis and the mean width of ten chondrocytes nearest to the symphysis. We found no significant differences in the mean chondrocyte width among pupfish species but observed increased chondrocyte density in untreated scale-eaters relative to generalists (Fig. 6e; Table S4; P = 0.009, ANOVA, Tukey’s HSD test). Moreover, only scale-eaters responded to M871, but not M35, by further increasing chondrocyte density relative to the control larvae (Fig. 6e; Table S4; P = 0.03, ANOVA, Tukey’s HSD test; mean ± 1 SE; control = 28.22 ± 0.38; M871 = 32.06 ± 1.06). Despite having shorter jaws after treatment with M871, chondrocyte density was significantly increased (Fig. 6e; mean ± 1 SE; control = 28.22 ± 0.38; M871 = 32.06 ± 1.06).

## Discussion

We used an evolutionary radiation of trophic specialist pupfishes, endemic to the Bahamas, to discover a novel function for *galr2a* in craniofacial divergence. Specifically, we confirmed that two transcription factor binding sites upstream of *galr2a* display highly divergent allele frequencies between trophic specialist species, visualized *galr2a* expression in craniofacial tissues in all three SSI pupfish species at two developmental stages and demonstrated a phenotypic effect on Meckel’s cartilage length and chondrocyte density using synthetic peptides to inhibit the activity of Galr2 and Galr1+2. Our findings demonstrate a crucial role for Galr2 in craniofacial divergence within this pupfish radiation. Our study also provides a roadmap in a non-model vertebrate system for rapidly identifying previously uncharacterized candidate genes important for adaptation to novel ecological niches (e.g., trophic specialization) that can be quickly validated through classic and state-of-the-art developmental biology tools. Overall, our research contributes to understanding the genetic basis of phenotypic evolution and adaptation in non-model organisms.

### Putative loss of a transcription factor binding site for *galr2a* in scale-eating pupfish

One of the most common evolutionary changes associated with phenotypic changes among closely related species is the gain or loss of cis-regulatory elements (Mack and Nachman, 2017; Signor and Nuzhdin, 2018). More than 85% of scale-eaters carry two transversions in the regulatory region of *galr2a* (Fig. 2). Combined with our observations of reduced *galr2a* expression in the mandibular mesenchyme during early jaw development in this species (Fig. 3), we conclude that the putative loss of a predicted *Sry* transcription factor binding site in scale-eaters is the most likely explanation for changes in gene expression, rather than a gain of a new predicted TFBS for *Znf416* at this locus (Fig. 2). This is further supported by the critical role of the *Sry*-related HMG box (Sox) family of transcription factors (especially the SoxE group including *Sox8, Sox9,* and *Sox10*) in craniofacial development as *Sry* transcription factors specify the behavior, multipotency, and survival of neural crest cells during vertebrate development (Leathers and Rogers, 2022; Simões-Costa and Bronner, 2015; Guo et al., 2015; Haldin and LaBonne, 2010; Betancur et al., 2010; Kamachi and Kondoh 2013).

Alternatively, we cannot rule out a gain of a TFBS upstream of *galr2a* in scale-eaters, additive or epistatic effects of both upstream transversions or more complex regulatory architectures, such as many interacting functional cis- and trans-acting regulatory variants or combinations of variants segregating at lower frequencies in trophic specialists that we have not prioritized (Boyle et al., 2017; McGirr and Martin, 2021; Price et al., 2015). However, our mapping cross of a single outbred pair of trophic specialists indicates that a single moderate-effect QTL containing *galr2a* explains 15% of phenotypic variation in oral jaw length between these species, consistent with causative variants affecting jaw size originating from this region (Martin et al. 2017).

### Differential *Galr2a* expression during development is associated with craniofacial divergence in SSI pupfishes

We found *galr2a* expression in our in situ hybridization experiments to be consistent with previously published RNAseq data for craniofacial tissue in this radiation (Fig. 3; Lencer and McCune, 2017; McGirr and Martin, 2018; McGirr and Martin, 2019; McGirr and Martin, 2021), with *galr2a* being differentially expressed only at 2 dpf. We observed distinctive spatial *galr2a* expression among SSI species at 2 and 8 dpf suggesting important time and tissue-specific regulation of *galr2a* expression during pupfish development. At 2 dpf, we found a strong association between decreased expression of *galr2a* in the mandibular mesenchyme anterior to the first pharyngeal arch with the future Meckel’s cartilage and oral jaw lengths in the adults of each species; with increasing *galr2* abundance in the molluscivores associated with shorter, but more robust jaws in adults (Martin and Wainwright 2013). In contrast, the reduced *galr2a* expression in the mandibular mesenchyme is associated with the development of longer oral jaws in scale-eaters, which is apparent as early as hatching time (Fig. 6d; Lencer et al., 2016).

We noted expression of *galr2a* in the maxilla of only molluscivores at hatching (Fig. 4 a-c), absent in the scale-eaters and generalists, consistent with the uniquely enlarged and anteriorly protruding head of the maxilla in this species (Martin and Wainwright 2013; Hernandez et al. 2018; Martin and Feinstein 2014; St. John et al. 2020). Furthermore, *galr2a* expression in the intermandibular muscles of the scale-eaters at hatching time (Fig. 4c) suggests that *galr2a* expression can also modulate the development of the observed hypertrophic musculature of the adductor mandibulae in the adult scale-eating pupfish (Hernandez et al., 2018). Altogether, these interspecific differences in spatial expression support a novel role for *galr2a* in musculoskeletal development and may contribute to the divergent craniofacial morphology observed in SSI pupfishes.

### Receptor inhibition supports a novel function for GALR2 in craniofacial divergence of SSI pupfishes

Interestingly, the response of Meckel’s cartilage length and chondrocyte density to the inhibition of Galr-receptor pathways was highly dependent on the species’ genetic background. Scale-eaters responded only to the Galr2-specific antagonist M871 with substantially reduced Meckel’s cartilage length and increased chondrocyte density but were unaffected by the Galr1+2 antagonist M35 (Fig. 6c). Importantly, M871 exhibits a higher binding constant for Galr2 receptors than the Galr1+2 antagonist M35 (Sollenberg et al., 2006; Sollenberg et al., 2010). Previous transcriptomic data from craniofacial tissues during early development (Lencer and McCune 2017) also indicates that *galanin* and *Galr1* mRNA transcript abundance does not vary among the three SSI species (Table S3).

Therefore, we conclude that even though scale-eaters expressed the lowest amount of *galr2a* transcripts and presumably contain overall fewer Galr2 receptors in their craniofacial tissue, M871’s higher binding affinity (Lang et al., 2007) was sufficient to inhibit these fewer receptors, resulting in reduced Meckel’s cartilage length (Fig. 7). In contrast, the presumably greater concentration of Galr2 receptors in molluscivore craniofacial tissue due to increased Galr2 expression during development resulted in the inhibition of these receptors by both M871 and M35, despite its lower binding affinity for Galr2 (Lang et al., 2007). We also conclude that inhibition of Galr1 receptors by M35 does not affect Meckel’s cartilage length or chondrocyte density and that the specific inhibition of Galr2 by M871 was sufficient to drive equal Meckel’s cartilage length shortening. Finally, the generalists’ lack of significant response to M35 is consistent with their decreased levels of *galr2a* at 2 dpf, suggesting that, as in scale-eaters treated with M35, fewer Galr2 receptors may be present for inhibition in these species’ background. However, the response to M871 appears to be specialist-specific, with little effect on the generalist Meckel’s cartilage length, and thus, we cannot rule out more complex species-specific factors interacting with Galr regulatory pathways.

**Figure 7.**
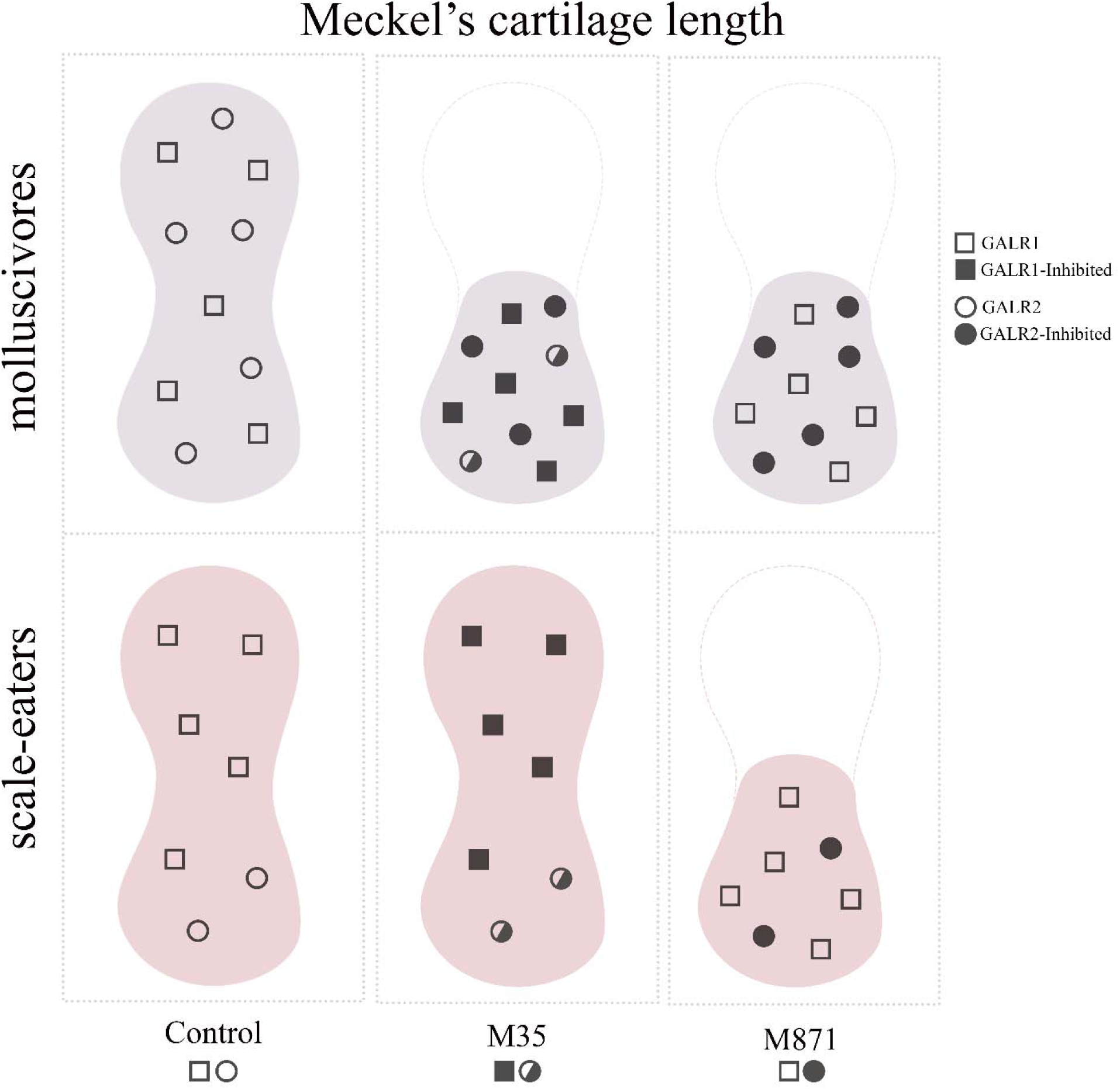
Proposed mechanism for how galanin receptor abundance affects Meckel’s cartilage length in the two pupfish trophic specialists after exposure to M35 (weak Galr1+Galr2 antagonist) or M871 (strong Galr2 antagonist). Molluscivores (top row) express more *galr2a* at 2 and 8 dpf than scale-eaters (bottom row) in their Meckel’s cartilage (see Figs. 3-4), resulting in more Galr2 receptors (open circles). Increased Galr2 receptor density in molluscivores results in their reduced Meckel’s cartilage length (see Fig. 6) due to inhibition by both the weak antagonist (M35: middle column; half-filled circles) and strong antagonist (M871: right column; filled circles) of Galr2. In contrast, reduced Galr2 receptor density in scale-eaters (bottom row) results in their reduced Meckel’s cartilage length only after exposure to the strong Galr2-specific antagonist M871.

Our work is consistent with previous work showing that *Galr2* activation inhibits cell proliferation in neuronal cell lines (Lang et al., 2007; Berger et al., 2004), suggesting that the TFBS allele frequency shifts observed in scale-eaters are the cause for low *galr2a* abundance and, as a consequence, have low Galr2 activity during craniofacial development resulting in increased chondrocyte density and Meckel’s cartilage length. Thus, the increased chondrocyte density observed in the M871-treated scale-eater larvae result likely from the strong and continued inhibition of the fewer Galr2 receptors available during development as a consequence or a synergy of/with their genetic background.

In conclusion, we propose that the inhibition of Galr2 and not Galr1 induces the reduction of Meckel’s cartilage length. Thus, a greater number of Galr2 receptors in the molluscivore specialist, as a result of increased expression of *galr2a* during early development, may result in their reduced oral jaw lengths as adults through increased opportunities for endogenous agonistic binding interactions (Fig. 7). Fewer Galr2 receptors on the scale-eater jaw may result in their enlarged jaw lengths as adults by limiting opportunities for Galr2 endogenous agonists to bind to these receptors during development. In the generalists, a greater number of Galr2 receptors but smaller overall Meckel’s cartilage length may limit the sensitivity of this species to Galr2 and Galr1+2 endogenous agonists, however, resulting in their intermediate length and least robust oral jaws among the three species under control conditions (Hernandez et al. 2018; Martin and Wainwright 2011).

### Conclusion

Our results support a novel role of the second receptor for galanin, Galr2, as a craniofacial modulator gene important in controlling craniofacial development and interspecific divergence through modifying its transcript abundance and receptor activity. *Galr2a* transcript abundance changes between species are associated with genetic changes in the regulatory region of *galr2a*, consistent with the loss of a transcription factor binding site in the scale-eating pupfish. We propose a model in which reduced Galr2 receptor density in the oral jaws of scale-eaters results in fewer endogenous agonistic interactions, increasing Meckel’s cartilage length presumably by decreased inhibition of chondrocyte proliferation (as a downstream effect of endogenous Galr2 activation). We also acknowledge the polygenic nature of jaw development and note that our previous genetic mapping experiments place an upper bound of 15% of oral jaw variation that may be due to differences in *galr2a* regulation among the myriad craniofacial genes driving the evolution of divergent craniofacial traits in this system.

## Materials and Methods

### Pupfish husbandry and maintenance

Three sympatric *Cyprinodon* species (*C. variegatus, C. brontotheroides,* and *C. desquamator*) were collected from both Osprey Lake and Crescent Pond in San Salvador Island, Bahamas in 2018 and maintained in the lab for at least two generations in small breeding groups. All lab colonies were maintained at a pH of 8.0 and 2-5 ppt salinity at 25-30°C on a 12:12 hours light:dark cycle at the University of California, Berkeley. Newly hatched fry were raised on brine shrimp nauplii for the first month and adults and juveniles were fed a combination of commercial pellet food and frozen bloodworms. All protocols and procedures employed were reviewed and approved by the University of California, Berkeley Animal Care and Use Committee (AUP-2015-01-7053) and animals were collected and exported from the Bahamas with research and export permits from the Bahamas Environmental, Science, and Technology commission through the Gerace Research Centre.

Multiple breeding pairs for each population were housed together in 40-liter aquaria with synthetic yarn spawning mops and fed ad libitum daily with frozen foods and commercial pellets. Embryos were collected by handpicking and raised at 27°C in 5 ppt salinity water with the addition of 10 mg/l gentamycin and 0.0001% methylene blue. Fertilization was confirmed after collection and embryos were staged according to Lencer and McCune (2018). The day of collection was noted as day “zero”, and thus one day after collection was 1-day post-fertilization (dpf).

### HCR FISH staining

Hybridization chain reaction (HCR) was used to visualize spatial expression of *galr2a* and *tpm3b* at key development timepoints (Choi et al., 2018; Ibarra-García-Padilla et al., 2021). HCR reagents, including probes, hairpins, and buffers, were purchased directly from Molecular Instruments (Los Angeles, California, USA). Staining was performed according to a modified protocol from Ibarra-García-Padilla et al., 2021, and Palominos and Whitlock, 2021.

#### Fixation and storage

2 and 8 dpf larvae were fixed in 4% paraformaldehyde (Electron Microscopy Science) 1X PBS (Invitrogen) overnight at 4°C. Samples were then serially dehydrated in methanol/1X PBS (25%, 50%, 75%) and stored in 100% methanol at −20°C for at least 16 hours before use.*<u>Permeabilization:</u>* Rehydration was performed serially in methanol/PBST (1X PBS 0.1% Tween20, Fisher Scientific) up to 100% PBST. Samples were then incubated for 10 minutes in pre-chilled acetone at −20°C for permeabilization. 2 and 8 dpf samples were then incubated for 20 and 80 minutes in 10 µg/ml Proteinase K (Thermofisher), respectively. <u>Prehybridization:</u> Samples were then pre-hybridized in the pre-warmed (at 37°C) hybridization buffer for 30 minutes at 37°C. *<u>Hybridization:</u>* For *Galr2a* and *Tpm3,* 4 to 6 pmol and 2 to 3 pmol were used, respectively. The probe mixture was pre-warmed for 30 minutes at 37°C before hybridization for 20 hours at 37°C. *<u>Washing probe excess:</u>* After hybridization, we performed four washes in pre-warmed wash buffer (15 minutes each) at 37°C, followed by two final washes in 5X SSCT (20X SSC from Thermofisher; SSCT is SSC with 0.1% Tween20) for 5 minutes each. *<u>Amplification:</u>* Preamplification involved incubating samples in pre-equilibrated (20-25°C) amplification buffer for 30 minutes. Next, amplification was performed using 4 µl of each snap-cooled hairpin in 250 µl of amplification buffer for 16 hours in the dark at room temperature. *Galr2a* expression was confirmed independently using two amplifiers with different excitation/emission wavelengths: B1-488 HCR amplifiers, and B1-594 HCR amplifiers. *Tpm3b* amplifiers used were B2-647 HCR amplifiers. *<u>Washing hairpin excess:</u>* Excess hairpins were washed several times with 5X SSCT, followed by a last wash with PBST. *<u>Nuclei labeling:</u>* After rinsing, samples were incubated for 30 minutes in DAPI/1X PBST (1 µm/ml, Sigma) and then washed in PBST 3 times for 5 minutes each. *<u>Clearing and mounting:</u>* Samples were then serially cleared in glycerol solutions at room temperature by sequentially removing and adding: 25% glycerol/75% PBST, 50% glycerol/50% PBST, 75% glycerol/25% PBST. Samples were then mounted in a custom-made bridged microscope slide (Fisher Scientific) in 75% glycerol/25% PBST.

#### Imaging

Samples were imaged using a Zeiss LSM880 (inverted) confocal microscope at the CNR Biological Imaging Facility at the University of California, Berkeley. All HCR were volumetric z-stack images of embryos or larvae positioned laterally or ventrally and taken every 3 µm. Images were deconvoluted, analyzed, and edited in FIJI (Schindelin et al., 2012) and Imaris (Bitplane).

### Galr2 inhibition during jaw development

According to the pupfish developmental series of Lencer and McCune (2018), the pharyngeal arches of *C. variegatus* are visible as segmented pouches ventral to the developing head at Stage 25 (48-64 hours post-fertilization at 27 °C, or 2 – 2.5 dpf), while the presence of a stomodeum-like oral cavity is evident by Stage 30 (100 – 130 hpf, or 4 – 5.5 dpf). We hypothesized that the key stages for proper jaw development occur between Stages 24 and 25, and thus we dechorionated Stage 24-25 embryos (or 2 to 2.5 dpf) pupfish by incubating them on 20 mg/ml Pronase (Roche) in water for 2.5 hours. The remaining chorions were removed manually with forceps. After larvae were dechorionated, we incubated them in 100 µM M35 (InnoPep Inc, San Diego, CA), a chimeric peptide antagonist to all galanin receptors (Galr1 and Galr2 in teleost fishes; Butler et al. 2020) and with a conserved 21 aminoacidic sequence to the core Galanin peptide sequence (GWTLNSAGYLLGPPPGFSPFR-NH2), or M871 (Abcam, USA), a Galr2-specific antagonist with the following aminoacidic sequence: WTLNSAGYLLGPEHPPPALALA-NH2, up to hatching time (Stage 33), looking for the following developmental events that are present in Stg 33 but not Stg 32: partial yolk absorption, frequent mouth movements with visible buccal pumping, frequent pectoral fin movements, and the ability to propel when touched. We also confirmed the formation of the ethmoid plate in both control and treated embryos after cartilage staining.

#### Cartilage staining

a non-acid clearing and staining method was used based on Walker and Kimmel, 2007, and Lencer and McCune, 2018. Fish larvae were cleared in 80% glycerol and mounted in custom-made coverslips (Fisher Scientific). Meckel’s length, inter-ocular distance, and chondrocyte density were quantified using the segmented line tool and ROI manager in Fiji (Schindelin et al., 2012). Inter-ocular distance was done by drawing a line from the center of one lens to the other and measuring the distance between the eyes (from the edge of the retinal pigmented epithelium). Imaging was done using a Zeiss Z1 AxioImager M2 microscope located at the Biological Imagining Facility of the University of California, Berkeley.

### Sequencing of *Galr2a* regulatory alleles

We used the DNeasy Blood and Tissue kit (Qiagen, Inc.) to extract DNA from the fins of wild-caught adult fish and quantified it on a Qubit 3.0 fluorometer (Thermofisher Scientific, Inc.). All specimens used in this study are kept in the Museum of Vertebrate Zoology at the University of California, Berkeley. The primers used to amplify both *Galr2a* regulatory loci are listed in Table S5. The sizes of the PCR products were confirmed by gel electrophoresis. PCR product clean-up was done using ExoSap-IT (USB) and quantified on a Nanodrop ND-1000 (Thermo Scientific) spectrophotometer. Sanger sequencing was performed at the UC Berkeley DNA Sequencing Facility. Sequence alignment was performed using Geneious Prime (Dotmatics).

### Prediction of transcription factors binding sites in *Galr2a-15kb* and *Galr2a-12kb*

We used the Multiple Expectation maximizations for Motif Elicitation (MEME) server (http://meme-suite.org/) and the motif database scanning algorithm TOMTOM (Bailey et al., 2009) to retrieve TFBS motifs from a 20kb conserved generalist, molluscivore, and scale-eater sequences upstream *galr2a* obtained from Richards et al., 2021. MEME suite finds, in the given sequences, the most statistically significant motifs first. We numbered the motifs following their statistical significance (Table S1).

### Statistics

#### Expression quantification of galr2a and tpm3b at 8 dpf

We segmented and quantified “jaw” *galr2a* expression in the maxilla, and the Meckel’s and palatoquadrate cartilages and “brain” *galr2a* expression in the central nervous system from HCR confocal images using 3D-Slicer (Fedorov et al., 2012). *Tpm3b* expression was segmented and quantified in the intermandibular muscles (IMA), the protractor hyoideus (PR-H), the adductor mandibular complex (AddM), and the levator arcus palatini (LEVs). Muscles were named according to Diogo et al., 2008. Statistical tests comparing the expression volume among species and tissues were performed in R (R Core Team 2023). We tested for significant differences in means via ANOVA and contrasted groups post-hoc using Tukey’s HSD test.

#### Expression quantification of garlr2a at 2 dpf

From the HCR confocal images, a total volume of 45 µm containing the pharyngeal arches was projected using the maximum intensity tool in Fiji. We selected two different tissues using ROI manager and quantified the relative fluorescence normalized by the background fluorescence of the image. First, we selected the mandibular mesenchyme plus the first and second pharyngeal arches (labeled in the figure “Pharyngeal Arches”). Then, we de-selected the second pharyngeal arch, quantifying only the expression of *galr2* in the mandibular mesenchyme and the first pharyngeal arch (labeled in the figure “Mandibular arch”). We then tested for significant differences in means via ANOVA and post-hoc analysis with Tukey’s HSD in R.

#### mRNA transcript analysis

A table containing RPKM values of transcripts from Lencer and McCune, 2017, datasets was obtained from the following Dryad repository, doi:10.5061/dryad.mb3gb. Genes of interest were found with the Gene ID listed on NCBI. Gene IDs are as follows: 106090243, 107092788, 107090827, 107096257, 107098818, and 107092130, for *galr2a, galr2b, galr1a, galr1b, gal,* and *sox21b*, respectively. Statistical tests comparing transcripts RPKM expression between species and developmental stages were performed in R, testing for significant differences in means via ANOVA and post-hoc analysis with Tukey’s HSD.

#### Cartilage staining

We compared chondrocyte density for significant differences in means using pairwise post-hoc Tukey’s HSD tests. We used a generalized linear model accounting for species and treatment and their interaction. We chose to measure chondrocytes within the Meckel’s cartilage because it allows for a clear view of each cell at the symphysis.

## Author’s contributions

Conceptualization: MFP, CTM, CHM; Data Collection: MFP, VM; Statistical analyses: MFP; Resources: CHM; Microscopy: MFP, VM; Original draft: MFP; Revising: MFP, EJR, CTM, CHM

## Acknowledgments

We thank Michelle St. John, David Tian, Mara Van Tassell, Chloe Clair, Dylan Chau, and Heidi Buratti for valuable comments and discussion of the results. We thank Lydia Smith for her help throughout the development of this project in the Evolutionary Genetics Lab at the University of California, Berkeley. We are particularly thankful to Austin H. Patton, who extracted the *C. brontotheroides* mRNA reads to create the HCR probes tested here, and for his insightful ideas and comments on the early stages of this manuscript. We thank the Gerace Research Centre and Troy Day for logistical support and the government of the Bahamas for permission to collect and export samples. This research was funded by the National Science Foundation DEB CAREER grant #1749764, National Institutes of Health grant 5R01DE027052-02, the University of North Carolina at Chapel Hill, and the University of California, Berkeley to CHM.

