## Supplemental Material for "Jaw size variation is associated with a novel craniofacial function for galanin receptor 2 in an adaptive radiation of pupfishes"

### Supplementary Methods

#### Alignment of the 20kb regulatory region with the *C. variegatus* genome

The two most differentiated SNPs upstream of galr2a were identified in a previous study (Richards et al. 2021). They occur at the following positions in the *Cyprinodon brontotheroides* genome assembly: c_bro_v1_scaf8:1003 and c_bro_v1_scaf8:4304 (GenBank: JAGKQL000000000.1, WGS: JAGKQL000000000.1-JAGKQL010015710) and the following positions in the *Cyprinodon variegatus* C_variegatus-1.0 genome assembly (GenBank: GCA_000732505.1): C_variegatus-1.0:858,678 and C_variegatus-1.0:862,008. One SNP in C_variegatus-1.0:858,678 is located at 11,310 base pairs upstream of the TSS of *galr2a* in *C. variegatus*. The second SNP found in the position C_variegatus-1.0:862,008 is located 7,980 base pairs upstream of the TSS of *galr2a* in *C. variegatus*.

### Supplementary Video S1

**Supplementary Video S1. *Galr2a* is expressed in the cytoplasm of developing chondrocytes in the Meckel's and Palatoquadrate cartilages.** A representative video of the jaw joint in an 8 days post-fertilization *C. variegatus* generalist pupfish. *Galr2a* is shown in orange, *tpm3b* in magenta, and nuclei are depicted in grey.

### Supplementary Figures S1-S5

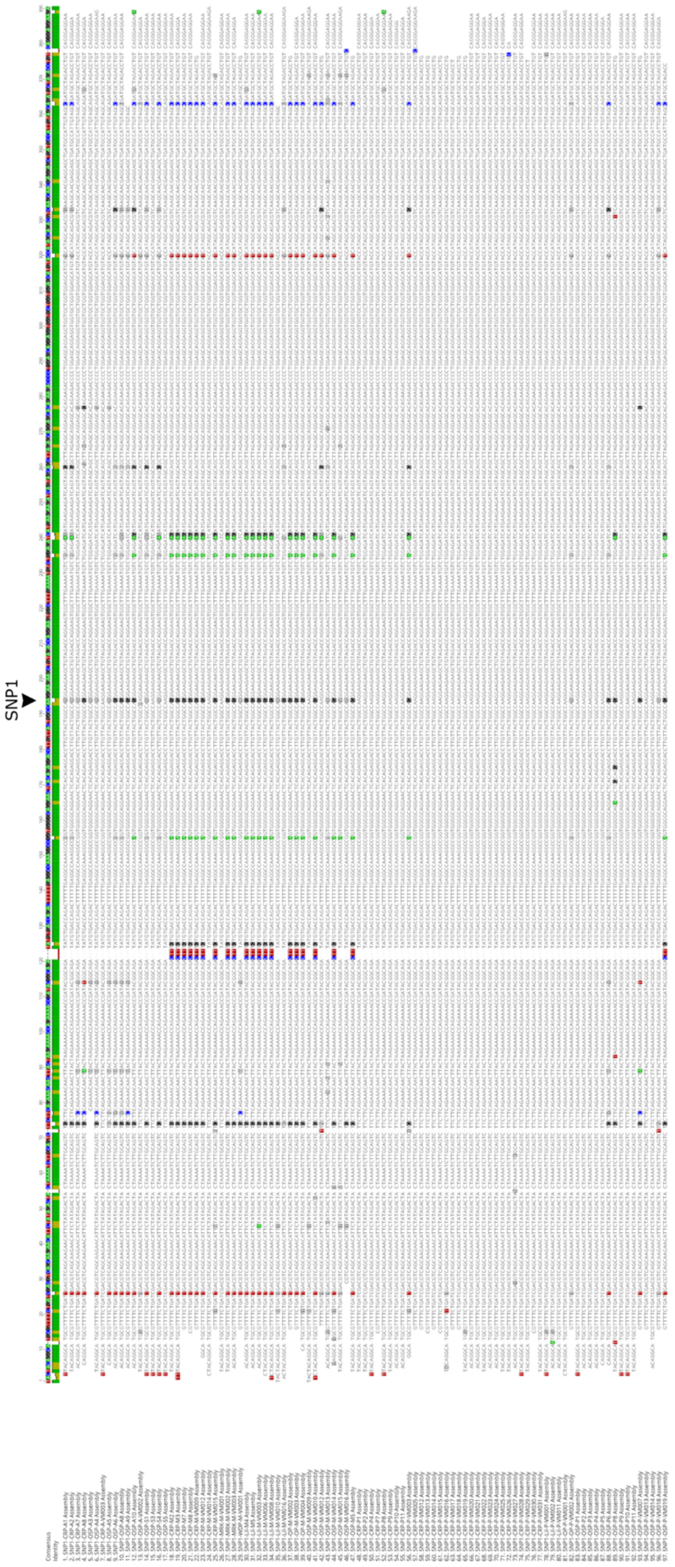
SNP1

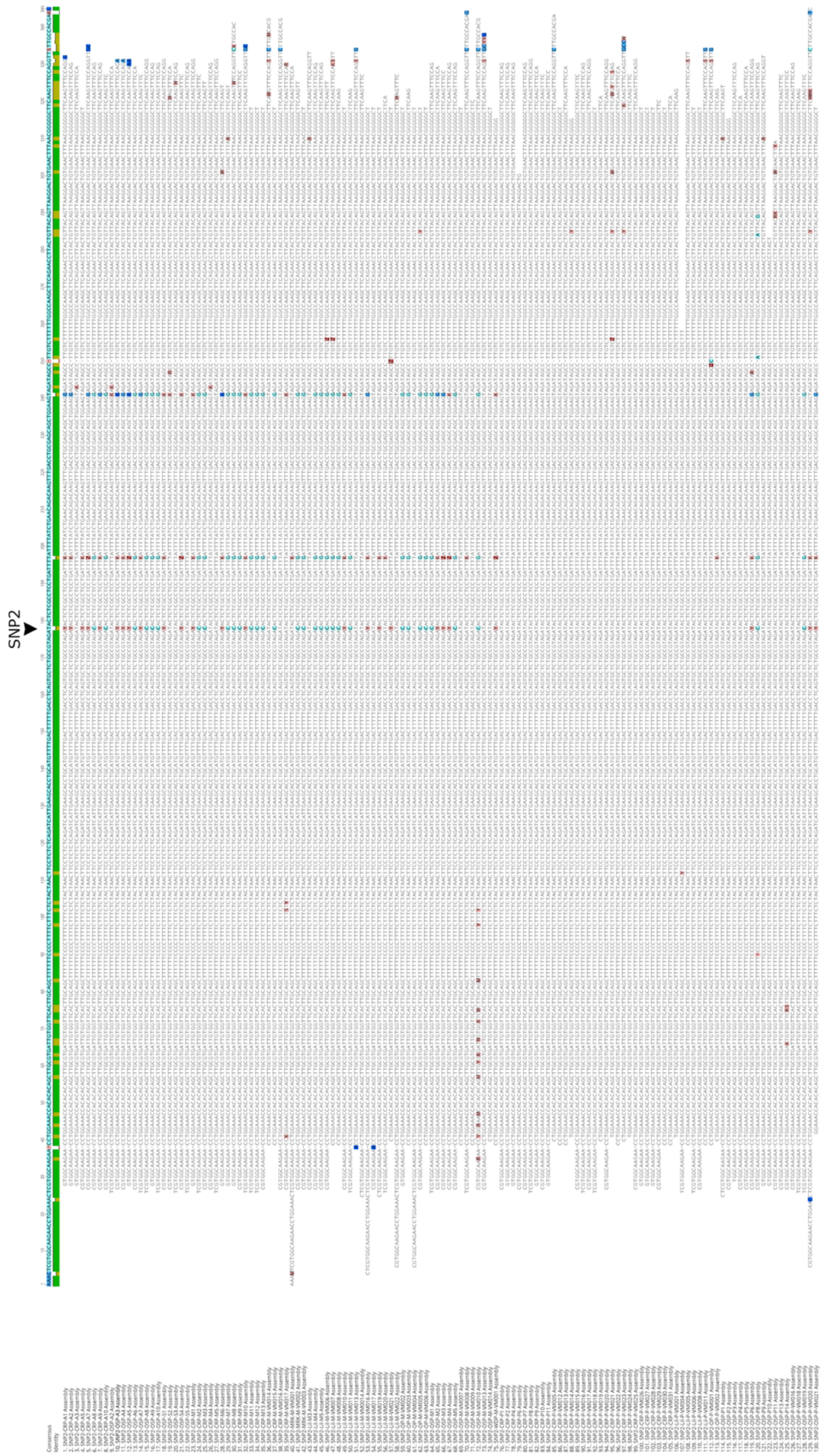


**Supplementary Figure S1 (page 3). Transversion from G/R in the generalist and molluscivore pupfishes to A in the scale-eater population.** The SNP was found in position 193

of the alignment and present in 86.3% of the total scale-eater individuals sequenced. CRP: Crescent Pond, Lil: Little Lake, MRK: Moon Rock Pond, OSP: Osprey Pond, OP: Ostrey Pond, WDP: Wild Dilly Pond.

**Supplementary Figure S2 (page 4). Transversion from C/Y in the generalist and molluscivore pupfishes to T in the scale-eater population.** The SNP was found in position 178 of the alignment and present in 90.9% of the total scale-eaters analyzed sequences and 100% of scale-eaters from Crescent Pond. CRP: Crescent Pond, Lil: Little Lake, MRK: Moon Rock Pond, OSP: Osprey Pond, OP: Ostrey Pond, WDP: Wild Dilly Pond.

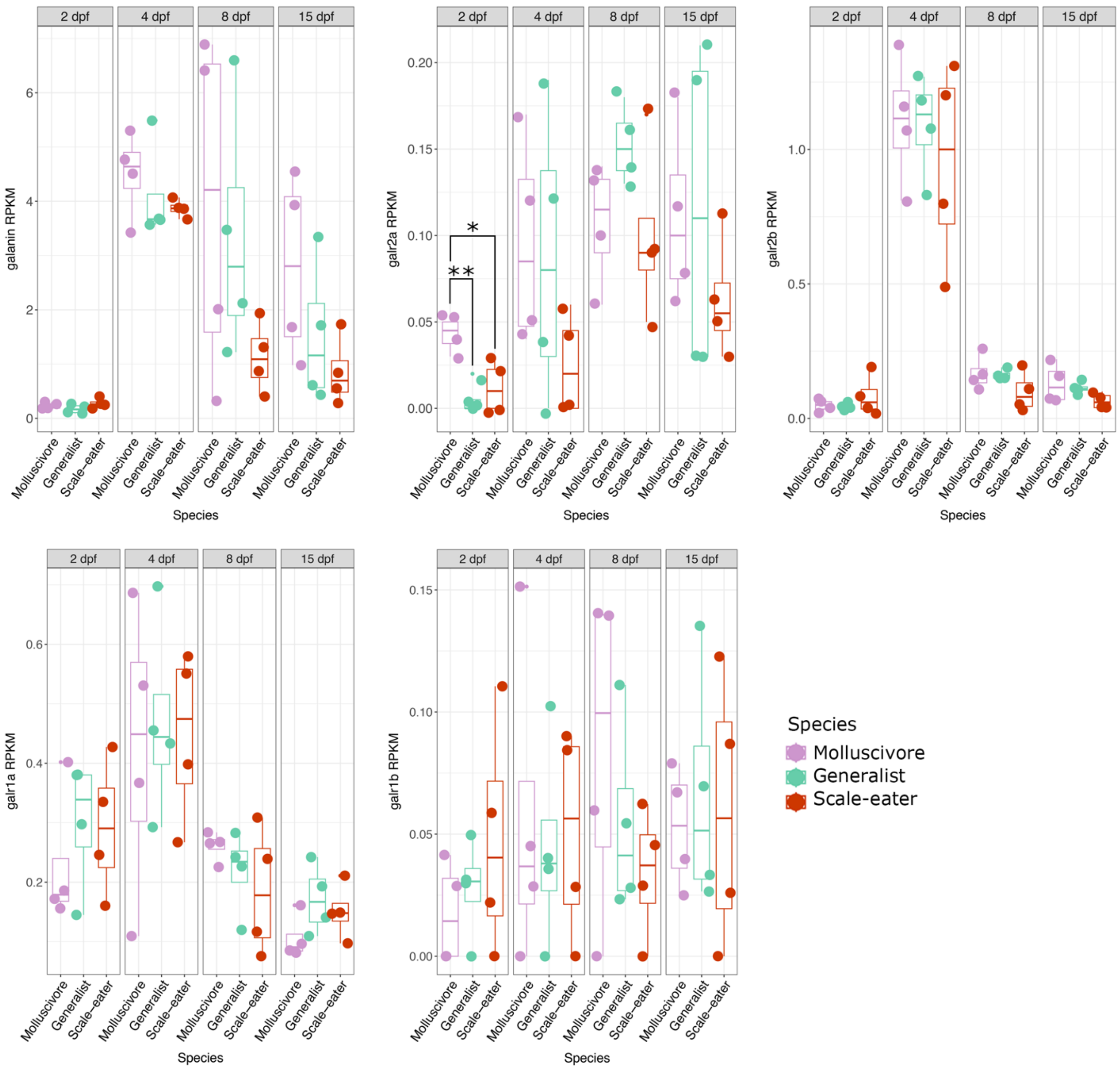


**Supplementary Figure S3. *galr2a* is the only transcript, in the galanin pathway, with different abundance at 2 days postfertilization between pupfish species.** Significant differences in means were tested via ANOVA and groups were compared post-hoc using Tukey's HSD test. Comparisons of *galanin*, *galr1a*, *galr1b*, and *galr2b* transcripts (RPKM) across species and developmental stages were not significant (P > 0.05), except for *galr2a*. The exact P-values for each comparison are listed in Table S2. Data from Lencer and McCune, 2017. $P < 0.05$, $^{**}P < 0.01$.

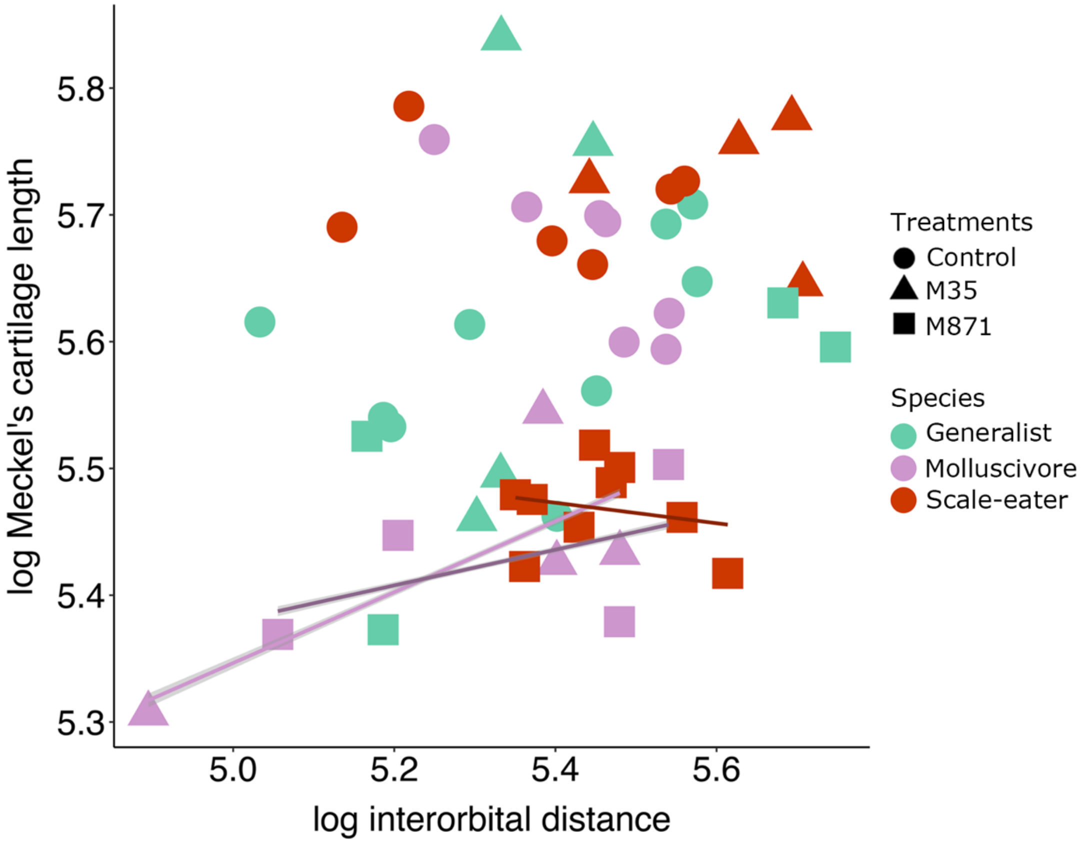


**Supplementary Figure S4. Allometric relationships between log-transformed Meckel's cartilage length and log-transformed interorbital distance across species show differences only in molluscivores and scale-eaters treated with M35 and M871.** Control (circle), M35-treated (triangle), and M871-treated (square). Darker regression lines indicate larvae that were treated with M871. No significant association was found between the Meckel's cartilage length and interorbital distance in control molluscivores, control and treated generalists, and M35-treated scale-eaters. Differences in Meckel's cartilage length and interorbital distance were found in M35- and M871-treated molluscivore larvae (P = 0.0006 and P = 0.04; linear regression model) and in M871-scale-eaters (P = 0.0095; linear regression model).

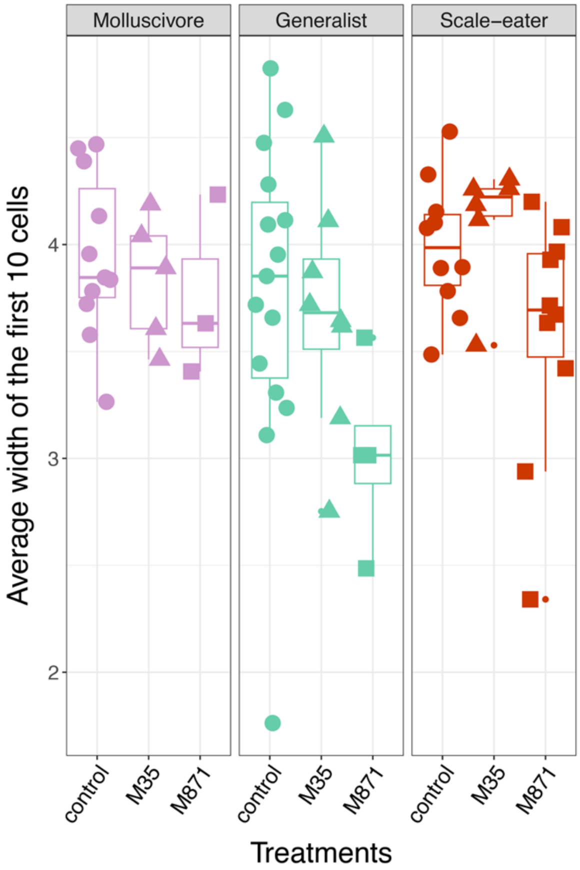


**Supplementary Figure S5. The average width of the first ten chondrocytes from the Meckel's cartilage symphysis does not change across species or treatments.** Significant differences in means were tested via ANOVA and groups were compared post-hoc using Tukey's HSD test. All comparisons were non-significant. The exact P-values for each comparison are listed in Table S4.

**Supplementary Tables S1-S5**

**Table S1. Predicted transcription factor bindings sites for the two single nuclear polymorphisms found in the 20kb *galr2a* regulatory region in the SSI pupfishes.**

| c_bro_b1_scaff8:1003 | Location from TSS | Sequence | p-value | ID | Tomtom Motif shared by TF | p-value |
|---|---|---|---|---|---|---|
| Generalists Molluscivores | -15,453 | CAGGgaA | 7.16E-04 | CAGGAA | Elf1 | 1.03E-04 |
| | | | | | Erg | 1.03E-04 |
| | | | | | GABPA | 1.41E-04 |
| Scale-eaters | | CAAGGAAga | 2.20E-04 | AGGASW | Elf5 | 6.35E-04 |
| | | | | | Spi1 | 7.84E-04 |
| | | | | | SPIB | 8.07E-04 |
| | | | | | Elf1 | 8.42E-04 |
| c_bro_b1_scaff8:4304 | | Sequence | p-value | ID | Tomtom Motif shared by TF | p-value |
| Generalists Molluscivores | -12,252 | ctgcAC | 2.17E-03 | MKRMAC | Sry_secondary | 3.39E-05 |
| | | | | | Sox21_secondary | 7.63E-04 |
| | | | | | Sox7_full | 2.08E-03 |
| Scale-eaters | | tAGATA | 7.70E-04 | YAGATA | ZNF416 | 1.73E-03 |

**Table S2. Differential expression of selected genes between pupfish species and developmental stages. Post-hoc Tukey's test analysis. Bolded values have a P-value < 0.05.**

| GeneID / Gene | Species Comparison | Stage (dpf) | P-value |
|---|---|---|---|
| **107090243 / Galr2a** | **Generalist (*C. variegatus*) x Molluscivore (*C. brontotheroides*)** | **2** | **3.81E-03** |
| 107090243 / Galr2a | Generalist (*C. variegatus*) x Molluscivore (*C. brontotheroides*) | 4 | 0.984 |
| 107090243 / Galr2a | Generalist (*C. variegatus*) x Molluscivore (*C. brontotheroides*) | 8 | 0.265 |
| 107090243 / Galr2a | Generalist (*C. variegatus*) x Molluscivore (*C. brontotheroides*) | 15 | 0.823 |
| 107090243 / Galr2a | Generalist (*C. variegatus*) x Scale-eater (*C. desquamator*) | 2 | 0.654 |
| 107090243 / Galr2a | Generalist (*C. variegatus*) x Scale-eater (*C. desquamator*) | 4 | 0.377 |
| 107090243 / Galr2a | Generalist (*C. variegatus*) x Scale-eater (*C. desquamator*) | 8 | 0.179 |
| 107090243 / Galr2a | Generalist (*C. variegatus*) x Scale-eater (*C. desquamator*) | 15 | 0.774 |
| **107090243 / Galr2a** | **Molluscivore (*C. brontotheroides*) x Scale-eater (*C. desquamator*)** | **2** | **0.014** |
| 107090243 / Galr2a | Molluscivore (*C. brontotheroides*) x Scale-eater (*C. desquamator*) | 4 | 0.304 |
| 107090243 / Galr2a | Molluscivore (*C. brontotheroides*) x Scale-eater (*C. desquamator*) | 8 | 0.958 |
| 107090243 / Galr2a | Molluscivore (*C. brontotheroides*) x Scale-eater (*C. desquamator*) | 15 | 0.364 |
| 107092788 / Galr2b | Generalist (*C. variegatus*) x Molluscivore (*C. brontotheroides*) | 2 | 0.987 |
| 107092788 / Galr2b | Generalist (*C. variegatus*) x Molluscivore (*C. brontotheroides*) | 4 | 0.996 |
| 107092788 / Galr2b | Generalist (*C. variegatus*) x Molluscivore (*C. brontotheroides*) | 8 | 0.992 |
| 107092788 / Galr2b | Generalist (*C. variegatus*) x Molluscivore (*C. brontotheroides*) | 15 | 0.892 |
| 107092788 / Galr2b | Generalist (*C. variegatus*) x Scale-eater (*C. desquamator*) | 2 | 0.47 |

| | | | |
|---|---|---|---|
| 107092788 / Galr2b | Generalist (*C. variegatus*) x Scale-eater (*C. desquamator*) | 4 | 0.766 |
| 107092788 / Galr2b | Generalist (*C. variegatus*) x Scale-eater (*C. desquamator*) | 8 | 0.31 |
| 107092788 / Galr2b | Generalist (*C. variegatus*) x Scale-eater (*C. desquamator*) | 15 | 0.427 |
| 107092788 / Galr2b | Molluscivore (*C. brontotheroides*) x Scale-eater (*C. desquamator*) | 2 | 0.554 |
| 107092788 / Galr2b | Molluscivore (*C. brontotheroides*) x Scale-eater (*C. desquamator*) | 4 | 0.715 |
| 107092788 / Galr2b | Molluscivore (*C. brontotheroides*) x Scale-eater (*C. desquamator*) | 8 | 0.264 |
| 107092788 / Galr2b | Molluscivore (*C. brontotheroides*) x Scale-eater (*C. desquamator*) | 15 | 0.182 |
| 107090827 / Galr1a | Generalist (*C. variegatus*) x Molluscivore (*C. brontotheroides*) | 2 | 0.65 |
| 107090827 / Galr1a | Generalist (*C. variegatus*) x Molluscivore (*C. brontotheroides*) | 4 | 0.93 |
| 107090827 / Galr1a | Generalist (*C. variegatus*) x Molluscivore (*C. brontotheroides*) | 8 | 0.71 |
| 107090827 / Galr1a | Generalist (*C. variegatus*) x Molluscivore (*C. brontotheroides*) | 15 | 0.06 |
| 107090827 / Galr1a | Generalist (*C. variegatus*) x Scale-eater (*C. desquamator*) | 2 | 0.99 |
| 107090827 / Galr1a | Generalist (*C. variegatus*) x Scale-eater (*C. desquamator*) | 4 | 0.987 |
| 107090827 / Galr1a | Generalist (*C. variegatus*) x Scale-eater (*C. desquamator*) | 8 | 0.815 |
| 107090827 / Galr1a | Generalist (*C. variegatus*) x Scale-eater (*C. desquamator*) | 15 | 0.444 |
| 107090827 / Galr1a | Molluscivore (*C. brontotheroides*) x Scale-eater (*C. desquamator*) | 2 | 0.72 |
| 107090827 / Galr1a | Molluscivore (*C. brontotheroides*) x Scale-eater (*C. desquamator*) | 4 | 0.98 |
| 107090827 / Galr1a | Molluscivore (*C. brontotheroides*) x Scale-eater (*C. desquamator*) | 8 | 0.37 |
| 107090827 / Galr1a | Molluscivore (*C. brontotheroides*) x Scale-eater (*C. desquamator*) | 15 | 0.27 |
| 107096257 / Galr1b | Generalist (*C. variegatus*) x Molluscivore (*C. brontotheroides*) | 2 | 0.899 |

| | | | |
|---|---|---|---|
| 107096257 / Galr1b | Generalist (*C. variegatus*) x Molluscivore (*C. brontotheroides*) | 4 | 0.94 |
| 107096257 / Galr1b | Generalist (*C. variegatus*) x Molluscivore (*C. brontotheroides*) | 8 | 0.65 |
| 107096257 / Galr1b | Generalist (*C. variegatus*) x Molluscivore (*C. brontotheroides*) | 15 | 0.94 |
| 107096257 / Galr1b | Generalist (*C. variegatus*) x Scale-eater (*C. desquamator*) | 2 | 0.67 |
| 107096257 / Galr1b | Generalist (*C. variegatus*) x Scale-eater (*C. desquamator*) | 4 | 0.98 |
| 107096257 / Galr1b | Generalist (*C. variegatus*) x Scale-eater (*C. desquamator*) | 8 | 0.83 |
| 107096257 / Galr1b | Generalist (*C. variegatus*) x Scale-eater (*C. desquamator*) | 15 | 0.95 |
| 107096257 / Galr1b | Molluscivore (*C. brontotheroides*) x Scale-eater (*C. desquamator*) | 2 | 0.42 |
| 107096257 / Galr1b | Molluscivore (*C. brontotheroides*) x Scale-eater (*C. desquamator*) | 4 | 0.98 |
| 107096257 / Galr1b | Molluscivore (*C. brontotheroides*) x Scale-eater (*C. desquamator*) | 8 | 0.34 |
| 107096257 / Galr1b | Molluscivore (*C. brontotheroides*) x Scale-eater (*C. desquamator*) | 15 | 0.99 |
| 107098818 / Galanin | Generalist (*C. variegatus*) x Molluscivore (*C. brontotheroides*) | 2 | 0.56 |
| 107098818 / Galanin | Generalist (*C. variegatus*) x Molluscivore (*C. brontotheroides*) | 4 | 0.71 |
| 107098818 / Galanin | Generalist (*C. variegatus*) x Molluscivore (*C. brontotheroides*) | 8 | 0.94 |
| 107098818 / Galanin | Generalist (*C. variegatus*) x Molluscivore (*C. brontotheroides*) | 15 | 0.12 |
| 107098818 / Galanin | Generalist (*C. variegatus*) x Scale-eater (*C. desquamator*) | 2 | 0.22 |
| 107098818 / Galanin | Generalist (*C. variegatus*) x Scale-eater (*C. desquamator*) | 4 | 0.89 |
| 107098818 / Galanin | Generalist (*C. variegatus*) x Scale-eater (*C. desquamator*) | 8 | 0.41 |
| 107098818 / Galanin | Generalist (*C. variegatus*) x Scale-eater (*C. desquamator*) | 15 | 0.97 |
| 107098818 / Galanin | Molluscivore (*C. brontotheroides*) x Scale-eater (*C. desquamator*) | 2 | 0.74 |

| | | | |
|---|---|---|---|
| 107098818 / Galanin | Molluscivore (*C. brontotheroides*) x Scale-eater (*C. desquamator*) | 4 | 0.45 |
| 107098818 / Galanin | Molluscivore (*C. brontotheroides*) x Scale-eater (*C. desquamator*) | 8 | 0.266 |
| 107098818 / Galanin | Molluscivore (*C. brontotheroides*) x Scale-eater (*C. desquamator*) | 15 | 0.06 |

**Table S3. Hybridization chain reaction expression differences in *galr2a* and *tpm3b* between species, tissues, and developmental stage. Tukey's Post-hoc analysis. Bolded values have a P-value < 0.05.**

| Species Comparison | Stage (dpf) | probe for | Tissue | P-value |
|---|---|---|---|---|
| Generalist (*C. variegatus*) x Molluscivore (*C. brontotheroides*) | 2 | *galr2a* | Pharyngeal Arches | 0.031 |
| Generalist (*C. variegatus*) x Molluscivore (*C. brontotheroides*) | 2 | *galr2a* | Mandibular Arch | 0.293 |
| Generalist (*C. variegatus*) x Scale-eater (*C. desquamator*) | 2 | *galr2a* | Pharyngeal Arches | 0.997 |
| Generalist (*C. variegatus*) x Scale-eater (*C. desquamator*) | 2 | *galr2a* | Mandibular Arch | 0.632 |
| Molluscivore (*C. brontotheroides*) x Scale-eater (*C. desquamator*) | 2 | *galr2a* | Pharyngeal Arches | 0.176 |
| Molluscivore (*C. brontotheroides*) x Scale-eater (*C. desquamator*) | 2 | *galr2a* | Mandibular Arch | 0.694 |
| Generalist (*C. variegatus*) x Molluscivore (*C. brontotheroides*) | 8 | *galr2a* | Brain | 0.745 |
| Generalist (*C. variegatus*) x Molluscivore (*C. brontotheroides*) | 8 | *galr2a* | Head | 0.228 |
| **Generalist (*C. variegatus*) x Molluscivore (*C. brontotheroides*)** | **8** | ***galr2a*** | **Jaw** | **0.0304** |
| Generalist (*C. variegatus*) x Scale-eater (*C. desquamator*) | 8 | *galr2a* | Brain | 1 |
| Generalist (*C. variegatus*) x Scale-eater (*C. desquamator*) | 8 | *galr2a* | Head | 0.554 |
| Generalist (*C. variegatus*) x Scale-eater (*C. desquamator*) | 8 | *galr2a* | Jaw | 0.212 |

| | | | | |
|---|---|---|---|---|
| Molluscivore (*C. brontotheroides*) x Scale-eater (*C. desquamator*) | 8 | *galr2a* | Brain | 0.727 |
| Molluscivore (*C. brontotheroides*) x Scale-eater (*C. desquamator*) | 8 | *galr2a* | Head | 0.768 |
| Molluscivore (*C. brontotheroides*) x Scale-eater (*C. desquamator*) | 8 | *galr2a* | Jaw | 0.502 |
| **Generalist (*C. variegatus*) x Molluscivore (*C. brontotheroides*)** | **8** | ***tpm3b*** | **Head** | **0.0418** |
| **Generalist (*C. variegatus*) x Scale-eater (*C. desquamator*)** | **8** | ***tpm3b*** | **Head** | **0.0282** |
| Molluscivore (*C. brontotheroides*) x Scale-eater (*C. desquamator*) | 8 | *tpm3b* | Head | 0.936 |

**Table S4. Meckel's cartilage length, chondrocyte density, and average chondrocyte width between species and treatments. Tukey's Post-hoc analysis. Bolded values have a P-value < 0.05.**

| Species | Measurement | Treatment comparison | P-value |
|---|---|---|---|
| Generalist (*C. variegatus*) | Meckel's length | Control - M35 | 0.994 |
| Generalist (*C. variegatus*) | Meckel's length | Control - M871 | 0.919 |
| Generalist (*C. variegatus*) | Meckel's length | M35 - M871 | 0.677 |
| **Molluscivore (*C. brontotheroides*)** | **Meckel's length** | **Control - M35** | **1.13E-03** |
| **Molluscivore (*C. brontotheroides*)** | **Meckel's length** | **Control - M871** | **9.47E-04** |
| Molluscivore (*C. brontotheroides*) | Meckel's length | M35 - M871 | 1 |
| Scale-eater (*C. desquamator*) | Meckel's length | Control - M35 | 1 |
| **Scale-eater (*C. desquamator*)** | **Meckel's length** | **Control - M871** | **6.34E-05** |
| **Scale-eater (*C. desquamator*)** | **Meckel's length** | **M35 - M871** | **2.03E-04** |
| Generalists vs. Scale-eaters | Meckel's length | Control - Control | 0.222 |
| Generalists vs. Molluscivores | Meckel's length | Control - Control | 0.749 |
| Scale-eaters vs. Molluscivores | Meckel's length | Control - Control | 0.99 |
| Generalist (*C. variegatus*) | Chondrocyte density | Control - M35 | 0.82 |
| Generalist (*C. variegatus*) | Chondrocyte density | Control - M871 | 0.578 |
| Generalist (*C. variegatus*) | Chondrocyte density | M35 - M871 | 1 |

| | | | |
|---|---|---|---|
| Molluscivore (*C. brontotheroides*) | Chondrocyte density | Control - M35 | 1 |
| Molluscivore (*C. brontotheroides*) | Chondrocyte density | Control - M871 | 1 |
| Molluscivore (*C. brontotheroides*) | Chondrocyte density | M35 - M871 | 1 |
| Scale-eater (*C. desquamator*) | Chondrocyte density | Control - M35 | 0.4 |
| **Scale-eater (*C. desquamator*)** | **Chondrocyte density** | **Control - M871** | **0.0322** |
| **Scale-eater (*C. desquamator*)** | **Chondrocyte density** | **M35 - M871** | **9.41E-05** |
| **Generalists vs. Scale-eaters** | **Chondrocyte density** | **Control - Control** | **9.26E-03** |
| Generalists vs. Molluscivores | Chondrocyte density | Control - Control | 0.112 |
| Scale-eaters vs. Molluscivores | Chondrocyte density | Control - Control | 0.965 |
| Generalist (*C. variegatus*) | Chondrocyte width | Control - M35 | 0.99 |
| Generalist (*C. variegatus*) | Chondrocyte width | Control - M871 | 0.22 |
| Generalist (*C. variegatus*) | Chondrocyte width | M35 - M871 | 0.505 |
| Molluscivore (*C. brontotheroides*) | Chondrocyte width | Control - M35 | 0.99 |
| Molluscivore (*C. brontotheroides*) | Chondrocyte width | Control - M871 | 0.99 |
| Molluscivore (*C. brontotheroides*) | Chondrocyte width | M35 - M871 | 0.99 |
| Scale-eater (*C. desquamator*) | Chondrocyte width | Control - M35 | 0.99 |
| Scale-eater (*C. desquamator*) | Chondrocyte width | Control - M871 | 0.72 |
| Scale-eater (*C. desquamator*) | Chondrocyte width | M35 - M871 | 0.589 |
| Generalists vs. Scale-eaters | Chondrocyte width | Control - Control | 0.97 |
| Generalists vs. Molluscivores | Chondrocyte width | Control - Control | 0.99 |
| Scale-eaters vs. Molluscivores | Chondrocyte width | Control - Control | 1 |

**Table S5. Primers used for Sanger sequencing the two identified single nuclear polymorphisms upstream *galr2a*.**

| Primers for | Forward | Reverse |
|---|---|---|
| galr2a-c_bro_b1_scaf8:1003 | ACAGGCATGCGTTTTCTGAT | TCCTCCCTGACAGGTCTAGC |
| galr2a-c_bro_b1_scaf8:4304 | CGTGGCAAGAACCTGGAAAC | GGAAACTTGAAGCCCCCGTTA |
